# Developmental adaptation of rod photoreceptor number via photoreception in melanopsin (OPN4) retinal ganglion cells

**DOI:** 10.1101/2023.08.24.554675

**Authors:** Shane P. D’Souza, Brian A. Upton, Kiara C. Eldred, Ian Glass, Kassidy Grover, Abdulla Ahmed, Minh-Thanh Ngyuen, Paul Gamlin, Richard A. Lang

## Abstract

Photoreception, a form of sensory experience, is essential for normal development of the mammalian visual system. Detecting photons during development is a prerequisite for visual system function - from vision’s first synapse at the cone pedicle and maturation of retinal vascular networks, to transcriptional establishment and maturation of cell types within the visual cortex. Consistent with this theme, we find that the lighting environment regulates developmental rod photoreceptor apoptosis via OPN4-expressing intrinsically photosensitive retinal ganglion cells (ipRGCs). Using a combination of genetics, sensory environment manipulations, and computational approaches, we establish a molecular pathway in which light-dependent glutamate release from ipRGCs is detected via a transiently expressed kainate receptor (GRIK3) in immature rods localized to the inner retina. Communication between ipRGCs and nascent inner retinal rods appears to be mediated by unusual hybrid neurites projecting from ipRGCs that sense light before eye-opening. These structures, previously referred to as outer retinal dendrites (ORDs), span the ipRGC-immature rod distance over the first postnatal week and contain the machinery for sensory detection (melanopsin, OPN4) and axonal/anterograde neurotransmitter release (Synaptophysin, and VGLUT2). Histological and computational assessment of human mid-gestation development reveal conservation of several hallmarks of an ipRGC-to-immature rod pathway, including displaced immature rods, transient *GRIK3* expression in the rod lineage, and the presence of ipRGCs with putative neurites projecting deep into the developing retina. Thus, this analysis defines a retinal retrograde signaling pathway that links the sensory environment to immature rods via ipRGC photoreceptors, allowing the visual system to adapt to distinct lighting environments priory to eye-opening.

---

Sensory experience plays a crucial role in development and establishment of the visual system^1,2^. Typically, visual experience is transmitted through conventional photoreceptors (rods and cones), to the remainder of the visual system, allowing it to use environmental features to shape its development^3^. However, with the recent discovery of atypical photopigments (Opn3, Opn4, and Opn5), and their widespread expression and functions^4^, it has become increasingly clear that non-image forming sensory experience plays a major role in development of the visual system and beyond^5^. In fact, several of these functions precede maturation of the conventional photoreceptors and conscious vision^5^. For example, prior to eye opening, OPN4 (melanopsin) and OPN5 (neuropsin) are each required for light-dependent vascular development of the retina and hyaloid artery^6,7^. In both cases, these photopigment-expressing retinal ganglion cells (RGCs) indirectly interact with immature vessel networks, sculpting the vascular landscape within the eye during development. In addition to developing vascular networks, OPN4-expressing intrinsically photosensitive retinal ganglion cells (ipRGCs) are thought to regulate inner retinal neuron numbers^6^, cone lamination^8^, retinal clock development^9^, and retinal wave maturation^10^. These findings suggest a fundamental and broad role for photoreception in development of the retinal landscape, prior to eye-opening.

In this current study, we expand on the roles of photoreception during development and find that ipRGCs regulate the number of rod photoreceptors in the immature retina. This occurs through a light-dependent process where ipRGCs promote the apoptosis of NRL+ immature rods restricted to the inner nuclear layer (INL) prior to eye-opening in mice. Additionally, using genetic and informatic approaches, we defined a mechanistic pathway linking glutamate release from ipRGCs to autonomous glutamate-detection via a transiently expressed *Grik3* kainate receptor in developing rods. Importantly, we identified putative communication structures on ipRGCs termed outer retinal processes, which emerge from the dendritic tree but molecularly resemble axons containing VGLUT2 and Synaptophysin. These structures project into the deep developing retina and interdigitate with immature rods, and possibly convey signals from ipRGCs. Finally, performing histological and informatic analysis on human retinal development, we find that fundamental features of this novel developmental pathway are conserved. Together, our results suggest that the neonatal retina tunes outer photoreceptor content to the environment through an atypical communication mechanism with ipRGC photoreceptors.

## Photoreception via OPN4 in ipRGCs promotes rod precursor cell death

OPN4 loss-of-function and dark reared mice show promiscuous retinal angiogenesis and a failure of hyaloid vessel regression thought to be caused by elevated retinal neuron number and increased oxygen demand^6^. To address the origin of this change, we compared the bulk retinal transcriptomes of postnatal day 3.5 (P3.5) *Opn4* null and wildtype mice reared in full-spectrum lighting (**Fig. 1a**, see Methods for details). Differential expression analysis revealed 255 genes that were melanopsin dependent (**Fig. 1b**). Using a Transfer learning approach (ProjectR)^11,12^, we mapped these differentially expressed genes to the single cell transcriptomes of developing retinal cells^13^ (**Fig. 1c**). This revealed two patterns (**Fig. 1d**) in which genes enriched in the *Opn4* null (**Fig. 1d**, Pattern 1) corresponded to developing rod photoreceptors (**Fig. 1c**, dark purple) and genes enriched in the wild type (**Fig. 1d**, Pattern 2) corresponded to retinal progenitor populations (**Fig. 1c**, orange, pink). Since these data suggested that melanopsin might regulate rod photoreceptor development, we deployed RNA Bisque^14^, a computational strategy that infers cell type proportions in bulk RNA-Seq by leveraging single cell transcriptomic data^13^. This showed that the proportion of rod photoreceptors was significantly increased (**Fig. 1e**) and the proportion of late retinal progenitors was significantly decreased (**Fig. 1e**, L-RPC) in *Opn4* null retina.

**Figure 1:**
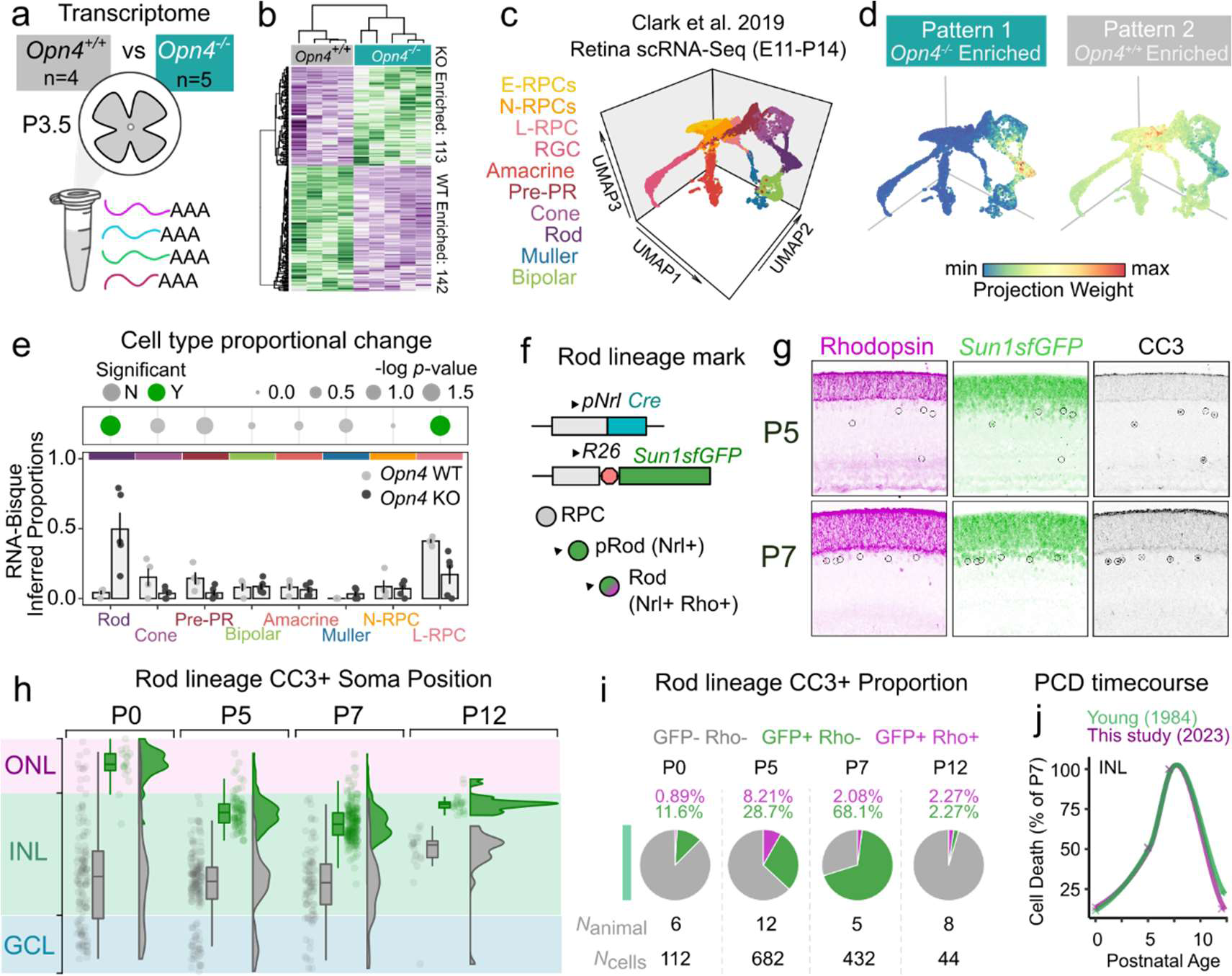
Transcriptomic profiling and spatial analysis of rod photoreceptors reveal a role for light and OPN4 signaling in rod apoptosis during development. **a-b**, Bulk RNA-seq profiling of pooled P3.5 *Opn4*^WT^ (*n =* 4 samples, 16 retina) and *Opn4*^KO^ littermates (*n =* 5 samples, 20 retina) identified 255 significantly differentially expressed genes (DEGs). **b**, Clustered heatmap representation of DEG expression (z-score) between WT and KO samples. **c.** 3D UMAP embedding of single cell transcriptomes from Clark et al., 2019 used for CoGAPS-assisted transcriptome mapping (see methods). **d**, Left, Projection weights of “Pattern 1” containing KO enriched genes (from **b**, left) and “Pattern 2” containing WT enriched genes (from **b**, right) onto cells from the atlas in **c**. **e**, Deconvolution and inference of the contribution from each cell class to the *Opn4*^WT^ and *Opn4*^KO^ transcriptomes using RNA-Bisque (2-way ANOVA, \*\*\*\**PCellClass × Genotype =* 8.81 × 10^−8^). Colored dot plot highlights significantly altered cell classes between WT and KO transcriptomes (\**P =* 0.0159, Wilcoxon test). **f-g**, Rod-INTACT lineage tracing employing the *Nrl-cre* transgene and cre-activated Sun1-sfGFP reporter line (*R26-* INTACT) to label immature rods (GFP+ Rho-) cells along their developmental trajectory in **g**. **h**, Spatial quantification of apoptotic cells (cleaved caspase 3+; CC3+) along the laminar axis of the retina, stratified by GFP expression. Data is represented as a triplex plot which includes individual soma points, boxplot, and normalized density estimates of laminar position for each age. **i**, Quantification of cell death in the INL attributable to immature rods (GFP+ Rho-), mature rods (GFP+ Rho+) or non-rods (GFP-Rho-) across development. GCL = ganglion cell layer, IPL = inner plexiform layer, INL = inner nuclear layer, NBL = neuroblastic layer, OPL = outer plexiform layer, ONL = outer nuclear layer. All bar graph data are presented as mean ± s.e.m.

A change in rod photoreceptor number in the *Opn4* null could be explained either by increased proliferation or decreased cell death within the lineage. Assessment of incorporation of the proliferation marker EdU at P4 and P9 and of cell division markers (pH3+ Ki67+ Vsx2+) did not reveal an effect of OPN4 loss of function (**Extended Data Fig. S1**). We then assessed the timing and pattern of cell death within the rod lineage. We used antibodies to cleaved caspase 3 (CC3) across a P0-P12 time course in the retinae of *Nrl-cre; Sun1sfGFP* mice (**Fig. 1f-h**) to map cell death timing and distribution. *Nrl-cre* is a rod lineage cre recombinase driver^15^ and *Sun1sfGFP* a nuclear envelope reporter that allows quantification of tightly packed cells^16^. Double labeling for GFP and rhodopsin (a marker of maturing rods) was used to identify rod developmental stage (**Fig. 1g**). Quantification of death for both GFP- and GFP+ (rod lineage) cells was mapped onto a model of retinal lamination (**Fig. 1h**) and, with rhodopsin labeling, presented as a proportion of the total dying cells (**Fig. 1i**). This showed that rod lineage cell death occurred primarily at the outer edge of the inner nuclear layer (INL)(**Fig. 1h**) and that at P7, was 71.18% of the total cell death in this layer (Fig. 1i). This demonstrates that rod lineage cells undergo programmed cell death, and identified the timing and localization of those events in retina experiencing a standard lighting environment. Notably, these data are consistent in timing and quantity with the original morphology-based description of retinal cell death from Young^17^ (**Fig. 1j**).

To determine whether rod lineage cell death was dependent on OPN4 and its photoreceptive activity, we assessed cell death in the retinae of *Opn4* null mice that were either reared in the dark (DD) or in standard lighting (LD)(**Fig. 2a**). Quantification of cell death at P7 (when rod lineage cell death is maximal (**Fig. 1f-i**) revealed that *Opn4* null mice showed a significantly reduced level (**Fig. 2b**, LD) that was restricted to the INL where immature rods reside. Furthermore, cell death in dark reared control, *Opn4^+/+^* mice (**Fig. 2b**, DD, gray bars) phenocopied the reduced level of cell death observed in the *Opn4* null under lighted conditions (**Fig. 2b**, LD, green bars). Notably, the level of cell death observed in DD conditions was unchanged by *Opn4* loss-of-function (compare **Fig. 2b**, DD, gray and cyan bars), suggesting that OPN4 is the only photopigment required for this response. Finally, we determined whether light-evoked apoptosis in the INL was specific to the rod lineage. Rearing *Nrl-cre; Sun1sfGFP* mice in complete darkness (**Fig. 2c**) led to decreased apoptosis in the INL (**Fig. 2d**). When dying cells were quantified for GFP expression, we observed a dramatic reduction in dying immature rods (GFP+) reared in darkness compared normally reared counterparts. This effect was limited to immature rods in the INL, as dying *Sun1sfGFP*-cells were equally abundant in both lighting conditions (**Fig. 2e**). Together, these data suggest that light, through melanopsin signaling, promotes the apoptosis of INL-localized immature rod photoreceptors.

**Figure 2:**
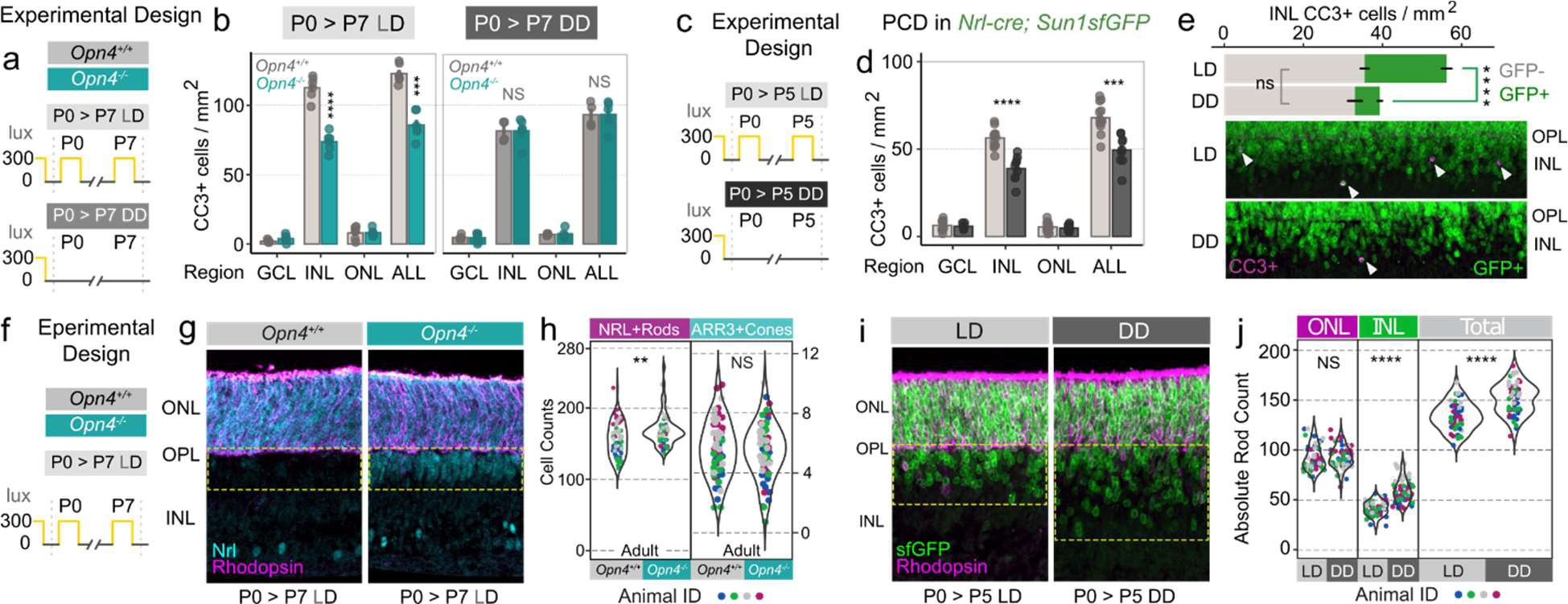
OPN4-signaling and light promote immature rod apoptosis and limit rod number over development. **a-b**, Dark-rearing paradigm comparing light-dark (LD) reared mice (*n* = 6 WT, *n =* 6 KO) and dark-reared (*n* = 4 WT, 5 KO) spatial apoptosis (Tukey HSD with fdr correction, \*\*\*\**P =* 5 × 10^−5^)**. c-d**, Dark-rearing paradigm comparing LD reared (*n* = 12 mice) or DD reared (*n* = 12 mice) P5 Rod-INTACT retinal apoptosis across region (\*\*\**P =* 7.72 × 10^−6^ INL, \*\*\**P =* 7.70 × 10^−5^ ALL; Tukey HSD test with fdr correction). **e**, Top, Quantification of immature rod and non-rod contribution of cell death in the INL in DD (*n* = 9 mice) and LD (*n =* 12 mice) *Nrl-cre; Sun1sfGFP* mice (\*\*\*\**P* = 3.52 × 10^−8^, Tukey HSD test with fdr correction). Bottom, Representative images of the dying rod band from LD and DD reared P5 Rod-INTACT mice. **f-g**, Analysis of maturing rods at P5 *Opn4*^WT^ and *Opn4*^KO^ litter mates reared in the cyclic light-dark cycle at P7, comparing Nrl+ Rho-(immature rods) and Nrl+ Rho+ (mature rods) reveals expansion of the INL immature rod population, consistent with dark-rearing experiments with *Nrl-cre; Sun1sfGFP* mice. **h,** Quantification of rod (Nrl+) and cone (Arr3+) numbers from adult *Opn4*^WT^ (*n =* 4 mice, 10,374 cells) and *Opn4*KO (*n* = 4 mice, 11,163 cells) littermates represented as absolute counts per field assessed, colored by animal (*\*\*P <* 0.01, Kolmogorov-Smirnov test). **i**, Representative images of LD and DD reared P5 *Nrlcre; Sun1sfGFP* mice stained for GFP and Rhodopsin. **j,** Analysis of rod numbers represented as or absolute counts per field (*n* = 4 per condition, \*\*\*\**P <* 0.0001, Kolmogorov-Smirnov test), colored by animal. GCL = ganglion cell layer, IPL = inner plexiform layer, INL = inner nuclear layer, NBL = neuroblastic layer, OPL = outer plexiform layer, ONL = outer nuclear layer. All bar graph data are presented as mean ± s.e.m.

The reduced rod lineage cell death in *Opn4* null and dark reared mice should result in increased numbers of rod photoreceptors. To assess this, we immunolabeled retinae from P7 *Opn4* null and control mice raised in standard lighting (**Fig. 2f**) with two rod lineage markers, NRL and Rhodopsin. This revealed an additional band of NRL+ cells at the outer edge of the INL in the mutant (**Fig. 2g**, yellow dashed boxes). When we quantified NRL+ rods and ARR3+ (arrestin) cones in adult mice, we identified a significant increase in the number of rods, but not cones (Fig. 2h). To assess the light dependence of this response, we raised *Nrl-cre; Sun1sfGFP* mice either in standard lighting (LD) or in complete darkness (DD) and quantified the number of GFP+ cells.

This showed an expansion of rod lineage cells in the INL of dark reared mice (**Fig. 2i**, yellow dashed boxes) that was significantly greater than controls reared in standard lighting (**Fig. 2j**). Combined, these data show, consistent with transcriptome analyses (**Fig. 1a-e**), that a light-melanopsin pathway normally promotes cell death within INL-localized immature rod photoreceptors, as a way of limiting the total number of mature rod photoreceptors.

## A transient ipRGC process extends into the outer retina and contains OPN4 and VGLUT2

ipRGCs and immature rods are widely separated in the developing retina. This raised the question of how ipRGCs transmit light information across the retina in a retrograde direction. In principle, ipRGCs could communicate *via* a long range soluble signaling ligand or over short range *via* a cellular process. For the former possibility we investigated a role for dopamine since it has been suggested that ipRGCs regulate levels of this neuromodulator as a way of controlling cone photoreceptor lamination^8^. However, when we generated mice with early retinal deletion of either *Th* (encoding Tyrosine Hydroxylase), or *Drd4* (encoding the dopamine receptor DRD4 expressed in the outer retina) we found that there was no consequence for patterns or quantity of cell death in the developing retina (**Extended Data Fig. S2**). We then considered the possibility that ipRGCs might use so called outer retinal dendrites (ORD)^8,18,19^ for retrograde communication. These processes were potential candidates for a communication conduit because they extend into the outer retina where rods and cone precursors reside^8^. As such, we analyzed their development and structure.

We used *Opn4^cre^* crossed to the cre-dependent reporter (*Ai14*) to mark ipRGCs. Upon recombination, *Ai14* produces cytoplasmic tdTomato fluorescent protein that allows visualization of neural processes. Between P0 and P7, this method exclusively labels ipRGCs within the inner retina (**Fig. 3a**, **Extended Data** Fig. 3a-c). Visualization of neural processes at higher magnification (**Fig. 3b**) revealed abundant fine processes in P0 retina that extended into the VSX2+ neuroblastic layer (NbL) of the outer retina. Quantification of outer retinal ipRGC processes over a developmental time-course revealed that they are transient and reduce in number by P5 and are shorter by P7 (**Fig. 3c, d**). These structures were not restricted to any given spatial territory in the retina (**Extended Data Fig. S3d**) and were not associated with dying ipRGCs (**Extended Data Fig. S3e**). In addition, an analysis of *Opn5* RGCs (**Extended Data Figure S3f**), which represent distinct RGC subtypes^20^, did not reveal any outer retinal processes, suggesting that they were exclusive to ipRGCs. Using the *Morf3* sparse lineage-tracing method^21^ (**Fig. 3f-h**, **Extended Data Fig. S4a**), we reconstructed whole ipRGCs to better understand the structure of outer retina processes. This showed that during early postnatal development (P0-P5) outer-stratifying ipRGCs resemble M1/M3 adult ipRGCs and project multiple outer retinal processes from their dendritic tree (**Fig 3f, g, Extended Data Fig. S4b**). As development proceeded, outer retina ipRGC processes became shorter until they were undetectable post eye-opening at P18 (**Fig 3h, Extended Data Fig S4c**). This is consistent with earlier analyses^18^.

**Figure 3:**
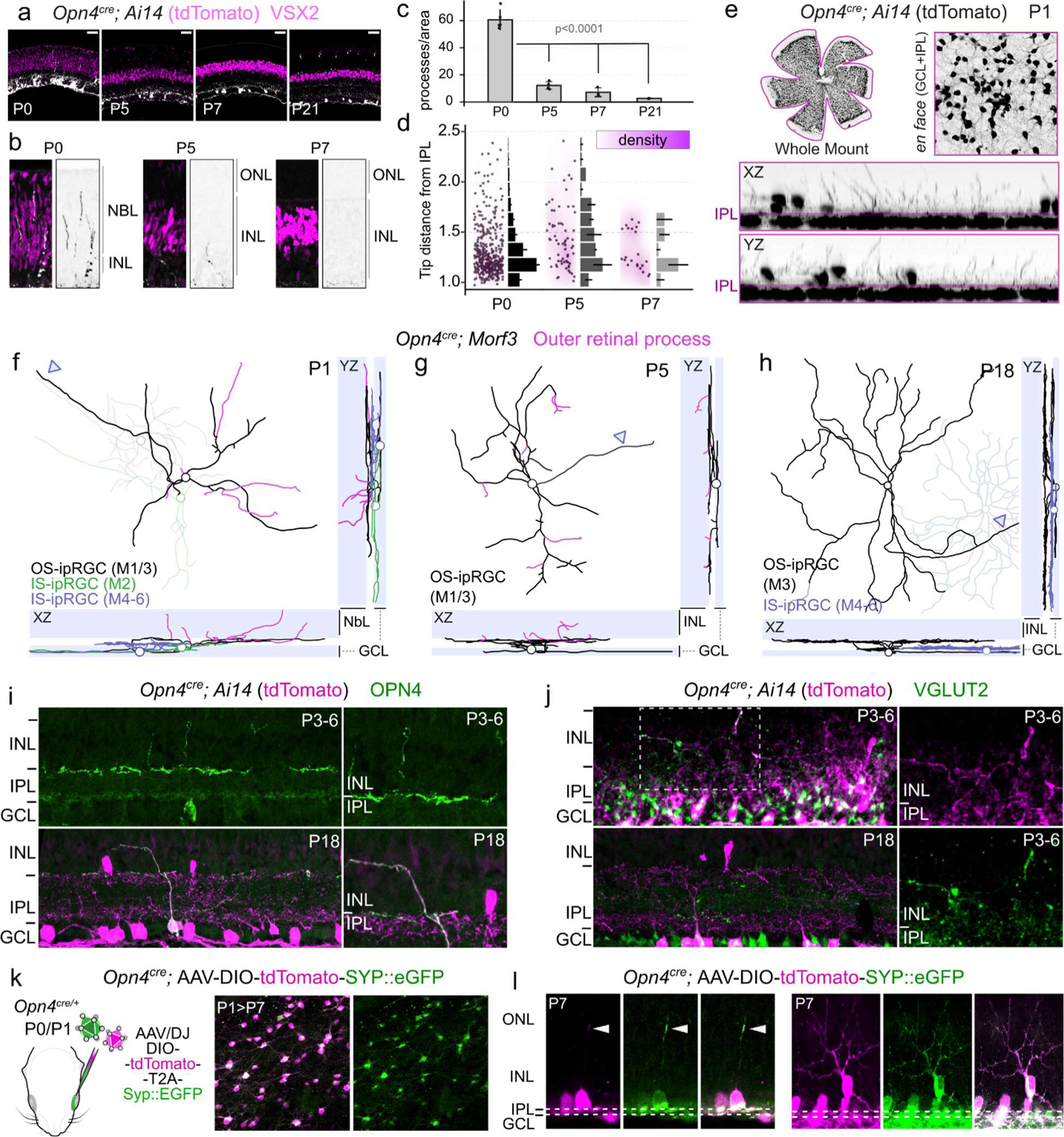
ipRGCs project hybrid neurites into the developing retina. **a-b**, Developmental retinal histology highlighting sprouts (grey, grey-inverted) and progenitor/bipolar cells (Vsx2, magenta). **c**, Quantification of sprout number (P0 *n* = 5; P5 *n* = 4, P7 *n* = 3, P21 *n* = 2, \*\*\*\*\**P =* 0.00005, One-Way ANOVA). **d**, Scaled process tip distance from IPL represented as a normalized density map and scaled histogram (P0: *n* = 380, 5 mice; P5: *n* = 69, 4 mice, P7: *n* = 22, 3 mice). **e**, Wholemount P1 *Opn4^cre^; Ai14* retina with orthogonal projection highlighting the depth of the retina and ipRGCs (grey-inverted). **e-h**, ipRGC reconstructions using the MORF3 sparse lineage mark (see S Fig 2 for details) representing cells from P1 (**e**), P5 (**f**), and P18 (**h**) projected en-face or orthogonally (XZ, YZ) for depth. Arrowhead refers to axon projecting towards the optic disc. **i-j**, Immunohistological labeling of sensory photopigment OPN4 (Melanopsin, **I**) and anterograde excitatory packaging proteins (Vglut2, **j**) in ipRGC sprouts over development (P3-6 to P18). **k**, Cre-dependent expression of bicistronic tdTomato and Synaptophysin-fused EGFP (Syp::EGFP) delivered to retina intravitreally at P0/1 and assessed at P7 (*n =* 29 mice). **l**, Representative spout-bearing ipRGCs expressing Syp::EGFP in dendrites during development. GCL = ganglion cell layer, IPL = inner plexiform layer, INL = inner nuclear layer, NBL = neuroblastic layer, OPL = outer plexiform layer, ONL = outer nuclear layer, RPE = retinal pigment epithelium, OS = outer stratifying cell, IS = inner stratifying cell. Data are presented as mean ± s.e.m.

Next, we evaluated the molecular properties of ipRGC processes. We found that outer retinal processes contain OPN4 immunoreactivity (**Fig. 3i**, **Extended Data Fig S5a**) regardless of developmental age^18,19^. However, mirroring their transience, we found that these processes accumulate VGLUT2 (the vesicular glutamate transporter that is necessary for RGC excitatory synaptic transmission)^22^, only during the postnatal period prior to eye-opening (**Fig 3j**, **Extended Data Fig. S5b**). These data suggest that the outer retinal processes of ipRGCs might be directly light sensitive, while the expression of VGLUT2 suggested that they might use the neurotransmitter glutamate to convey light information to the outer retina.

The transmission of information in the retrograde direction within the retina would be unusual and an activity more commonly associated with axons, not dendrites. To clarify the nature of these processes, we performed two types of labeling for synaptophysin (SYP), a presynaptic marker. Labeling of *Opn4^cre^; Ai14* marked ipRGCs with antibodies to SYP identified outer retinal processes coincident with tdTomato (**Extended Data Fig. S5c**). To overcome the limitation arising from the high density of SYP within the retina, we also used cre-dependent viral delivery of SYP fused to eGFP (SYP::EGFP) in *Opn4^cre^* mice (**Fig. 3k, l**, **Extended Data Fig. S5d, e**). This resulted in robust accumulation of this pre-synaptic vesicular docking protein in outer retinal processes (Fig. 3l, **Extended Data Fig. S5d**). As observed with VGLUT2 accumulation in retinal processes during development, SYP accumulation in ipRGC dendrites was observed at P7 (**Extended Data Fig S5d**) but was greatly diminished at P28 (**Extended Data Fig. S5e**) suggesting a developmental regulation of its distribution. These data suggest that outer retinal ipRGC projections are unusual hybrid neurites with anatomical origins of dendrites but molecular characteristics of axons.

## ipRGCs use a light-OPN4-glutamate pathway to induce rod precursor cell death

Given the evidence suggesting ipRGCs may directly communicate with immature rods and regulate their apoptosis, we set out to determine whether process-bearing ipRGCs were capable of detecting light early in the first postnatal week. We stimulated day-of-birth dark-adapted *Opn4^cre^; Ai14* pups with 470 nm light to activate ipRGCs and assessed activity using cFos as an immediate early gene (IEG) read-out (**Fig. 4a**). Pups exposed to light showed a dramatic increase in the number of cFos+ cells in the GCL and INL compared with dark-adapted littermates (**Fig. 4b**). cFos induction was only observed in tdTomato+ cells from light-stimulated animals (**Fig. 4c, d**). Furthermore, many of the cFos+ ipRGCs projected neurites into the outer retina (**Fig. 4e, f**). In the brain, significant increases in cFos+ cells were observed within the thalamic intergeniculate leaflet (IGL), a region known to receive input from outer-stratifying M1 ipRGCs (**Fig. 4g-i**)^23,24^. These data indicate that ipRGCs that stratify to the outer retina are photosensitive from the day of birth.

**Figure 4:**
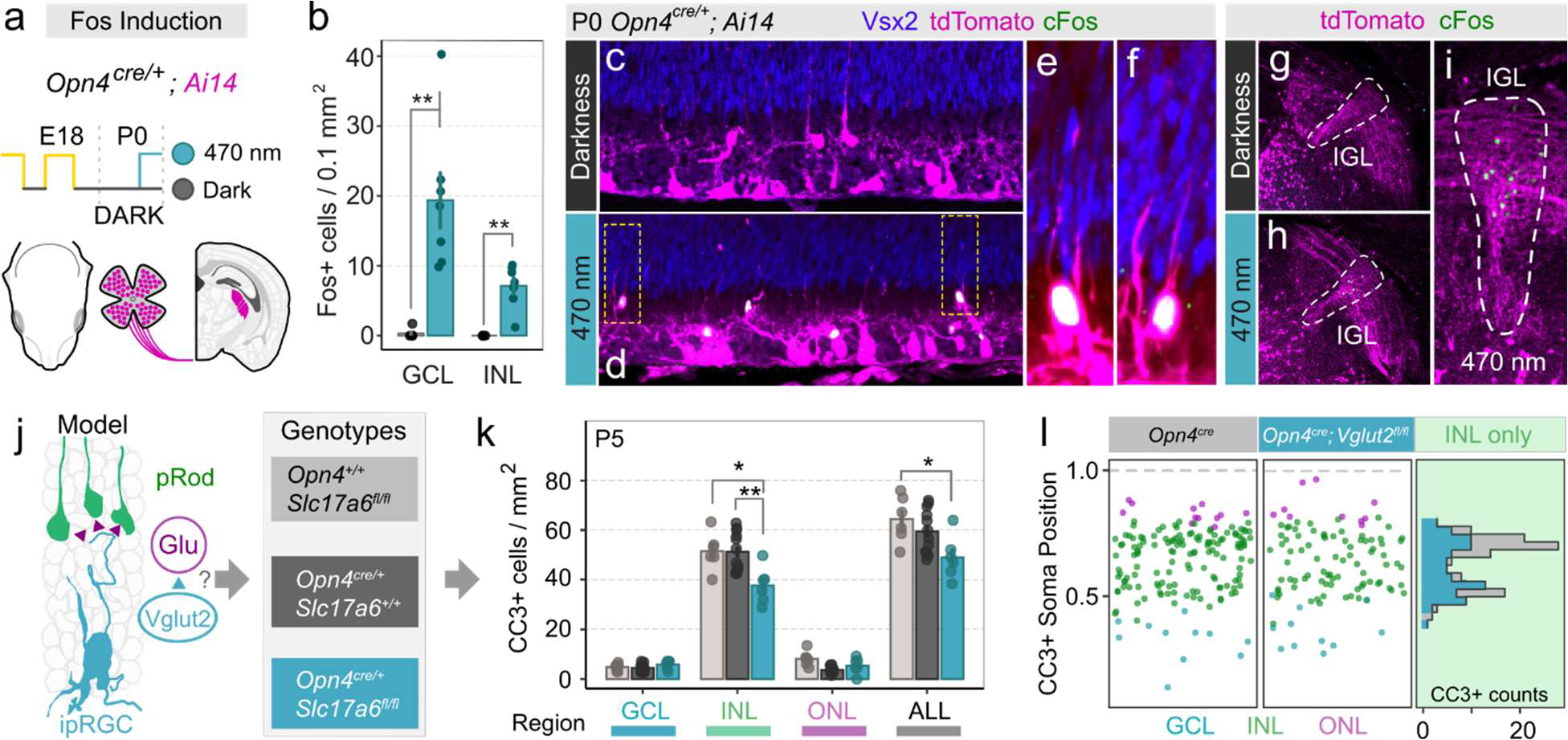
Glutamate-release from ipRGCs regulates immature rod apoptosis. **a**, Fos-activation paradigm in *Opn4^cre/+^; Ai14* mice stimulated with 1 × 10^14^ photons cm^−2^ sec^−1^ 470 nm light for 90 minutes at P0. **b**, Quantification of Fos-induction in the GCL and INL of dark-adapted (black circles; *n =* 5 mice) or light-stimulated (cyan circles, *n* = 7 mice) mice (\*\**P <* 0.005, Wilcoxon test with fdr correction). **c-f**, Representative images highlighting ipRGCs (magenta), progenitor cells (blue, VSX2+), and Fos (green) in the GCL and INL. **e-f**, Magnification of Fos+ ipRGCs from retina of light-stimulated mice highlight multiple deep projecting sprouts. **g-i**, Thalamic IGL Fos induction (green) in dark adapted and light-stimulated mice. **j**, Schematic model of direct glutamate release from sprout-bearing ipRGCs. **k**, Assessment of regional apoptosis in control (*Slc17a6^fl/fl^, n =* 6 mice) or *Opn4^cre/+^*(*n =* 12 mice) and Vglut2-cKO (*Opn4^cre/+^; Slc17a6^fl/fl^, n =* 6 mice) retina at P5 (\**P =* 0.014 control vs. Vglut2-cKO INL; \*\**P* = 0.005 *Opn4^cre/+^* vs. Vglut2-cKO INL, \**P =* 0.018 control vs Vglut2-cKO Total, fdr corrected Tukey HSD between genotypes per region). **l**, Spatial plot of CC3+ soma positions from control and Vglut2-cKO P5 retina. GCL = ganglion cell layer, IPL = inner plexiform layer, INL = inner nuclear layer, NBL = neuroblastic layer, OPL = outer plexiform layer, ONL = outer nuclear layer, RPE = retinal pigment epithelium, OS = outer stratifying cell, IS = inner stratifying cell. Data are presented as mean ± s.e.m.

In adulthood, ipRGCs release multiple classes of neurotransmitter including glutamate (excitatory)^25,26^, PACAP (modulatory)^26^ and GABA (inhibitory)^27^ to influence circadian entrainment and the pupillary light reflex. Using a scRNA-Seq atlas of developing RGCs over fetal and early postnatal development^28^, we identified developing M1s (see Methods for details) and assessed expression of genes related to neurotransmitters or their packaging (**Extended Data Fig. S6a-c**). From E14-P5, M1 ipRGCs consistently express melanopsin (*Opn4*) and VGlut2 (*Slc17a6*). A subset also express PACAP (*Adcyap1*) and Gad2 (*Gad2*). Given that VGlut2 accumulates in ipRGC sprouts during early postnatal development (**Fig. 3**), we assessed whether release of glutamate was a necessary component of light-evoked immature rod apoptosis.

Using an *Slc17a6 loxp* flanked allele (*Slc17a6^fl^*)^29^ and *Opn4^cre^*, we generated a VGLUT2 loss-of-function in ipRGCs (Fig. 4j)^25,26^. Conditional deletion of *Slc17a6* did not change the number of ipRGCs that developed (**Extended Data Fig. S6d**) nor the number of outer retina processes (**Extended Data Fig. S6d, e**). A significant reduction in the length of outer retinal processes in VGLUT2 loss-of-function ipRGCs (**Extended Data Fig. S6f, g**) was observed, at P0 (12 μm) and P5 (20.2 μm), but was small in absolute terms. Furthermore, *Opn4^cre^*; *Slc17a6^fl/fl^* mice showed normal light-induced cFos in ipRGCs (**Extended Data Fig. S6h, i**). However, as expected given the established role of glutamate in ipRGC signaling, postsynaptic light-induced cFos in the suprachiasmatic nucleus (SCN) was dramatically diminished (**Extended Data Fig. S6j**). These data indicate that VGLUT2 loss-of-function did not compromise the proximal mechanisms of photoreception^25^ but did compromise downstream glutamate signaling.

Assessment of cell death in the retina of *Opn4^cre^*; *Slc17a6^fl/fl^* revealed a significant reduction compared to control littermates (**Fig. 4k**). Importantly, when cell death events were mapped spatially (**Fig. 4l**) reduced cell death was restricted to the outer INL, consistent with the positioning of immature rod photoreceptors (**Fig. 1h**). Combined, these analyses indicate that the death of immature rods is dependent on light-dependent glutamate signaling.

## GRIK3 is an immature rod glutamate receptor required for cell death

The proposed mechanism for ipRGC induction of rod precursor cell death (**Fig. 5a**) requires that rod precursors express a glutamate receptor. Reanalysis of the Clark et al dataset^13^ using Monacle3^30^, defined rod lineage developmental progression (**Fig. 5b**) and a pseudotime axis for that lineage (**Fig. 5c**). Assessment of glutamate receptor expression within this lineage revealed that most were not expressed (data not shown). Among those that were expressed within the rod lineage, only *Grik3*, a low-affinity kainate receptor, appeared to be an ideal candidate based on its expression and temporal pattern. Mapping *Grik3* expression onto the rod development pseudotime axis showed that it was transiently expressed with a timing that matched very closely the rise in expression of *Nrl*, the crucial rod lineage transcription factor (**Fig. 5d, e**). The timing of transient *Grik3* transcript expression was confirmed by assessing bulk transcriptome data from developmentally staged, flow sorted rod lineage cells marked with *Nrl-eGFP*^31^. This showed that *Grik3* transcript was high between P2-P6 and diminished thereafter (**Fig. 5f**). Furthermore, labeling of a P0-P12 developmental retina series with an antibody to GRIK3 showed similar expression transience and a distribution that emphasized the outer edge of the INL (as marked by *Nrl-cre; Sun1sfGFP*; **Fig. 5g**), the precise location to which ipRGC processes stratify and light-regulated rod apoptosis occurs. Together, these data identified GRIK3 as a putative receptor in glutamate-dependent death of developing rods.

**Figure 5:**
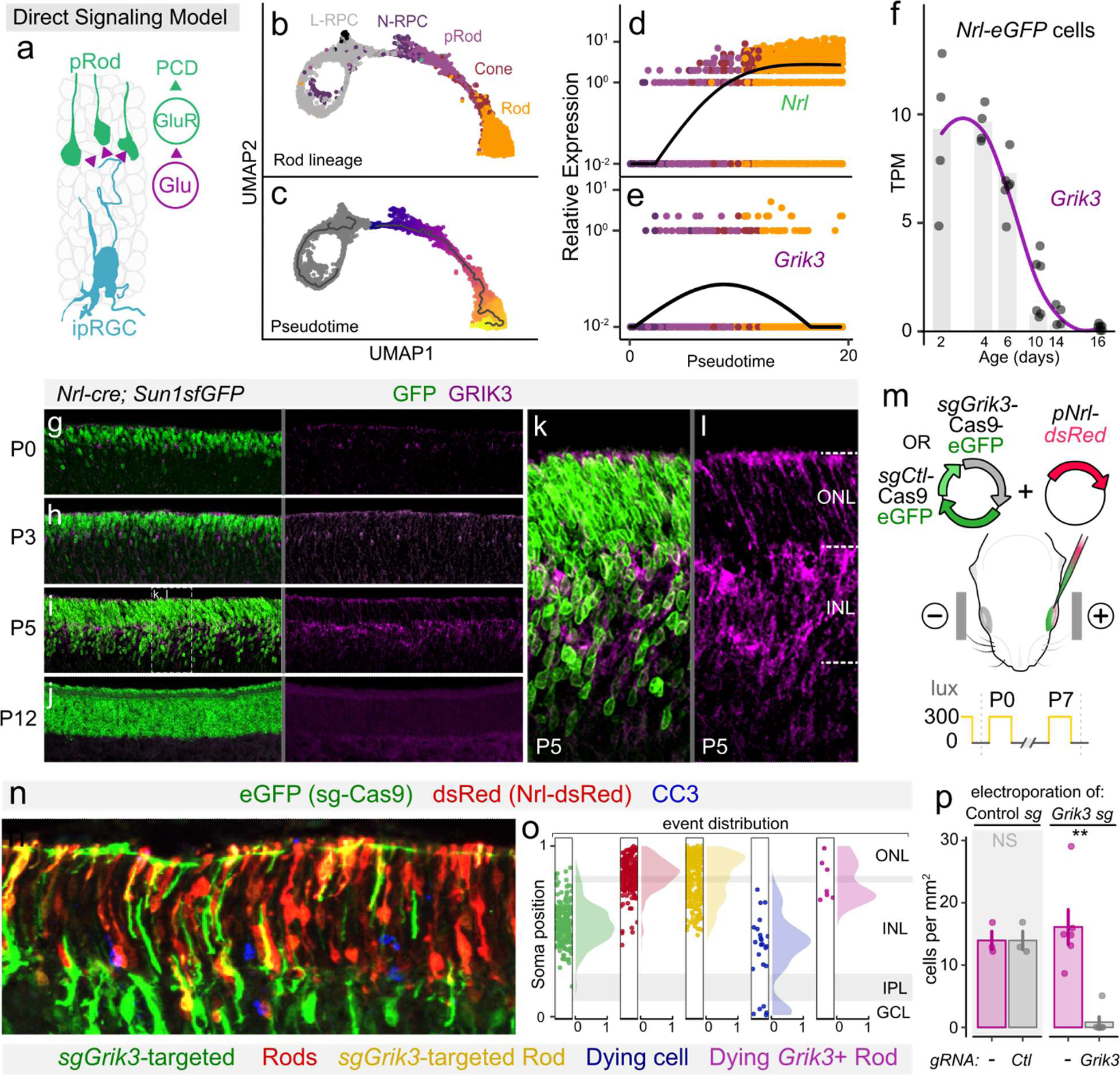
Transiently expressed GRIK3 regulates rod apoptosis during development. **A**, Schematic model of direct glutamate detection via immature rods, via an unknown GluR. **B-e**, UMAP embedding and pseudotime analysis of rod-lineage from Clark et al., 2019 reveals transient expression of *Grik3*. **F**, Confirmatory analysis of *Grik3* expression dynamics over postnatal development from Kim et al., 2016 which performed bulk transcriptomic assessment of immature rods flow sorted from *Nrl-eGFP* retina. **G-j**, Time course evaluation of Grik3 protein expression *in vivo,* using *Nrl-cre; Sun1sfGFP* to lineage-mark developing rods. **K-l**, Magnification of INL-ONL border highlighting expression of GRIK3 in INL-developing rods. **M**, *In vivo* electroporation strategy to somatically knockout *Grik3* from the developing retina (*sgGrik3*-Cas9) while simultaneously labeling postnatally born rods (*pNrl-*dsRed). **N**, Histological confirmation at P7 of high-efficiency electroporation in the outer retina labeling transfected (dsRed+ or eGFP+) and dying cells (CC3+). **O,** Spatial localization and scaled density plots of 665 cells from *n =* 6 mice co-electroporated as described in **m-n**. **p,** Analysis of cell death of GFP+ (grey) and GFP-(magenta) electroporated rod population (dsRed+) between Control-*sg* and *Grik3*-sg electroporated mice (*n =* 3 Control, *n =* 6 *Grik3-*sg, \*\**P =* 0.0037, Wilcox test between Cas9 constructs). GCL = ganglion cell layer, IPL = inner plexiform layer, INL = inner nuclear layer, NBL = neuroblastic layer, OPL = outer plexiform layer, ONL = outer nuclear layer. Data is represented as mean ± s.e.m.

To assess GRIK3 function, we used somatic CRISPR gene targeting. P1 mice were electroporated in the retina^32,33^ with *Nrl-dsRed*^34^ (to mark rod lineage cells) and with *sgGrik3-Cas9-EGFP* encoding guide RNAs (sgRNAs) against exon 4 of *Grik3* and eGFP for cell marking (**Fig. 5m**). A control *sgCtl-Cas9-eGFP* plasmid omitted *Grik3* guide RNA sequences. Electroporation at P1 produced high-efficiency expression of both *Nrl-dsRed* and *sgGrik3-Cas9-EGFP* plasmids (**Fig. 5n**) at P7. After harvesting at this developmental stage, retinas were labeled for cleaved caspase 3 (in blue) to identify dying cells. Importantly, when fluorescent protein expression was mapped onto a model of retinal layers (**Fig. 5o**) they were distributed as would be expected with *eGFP*, driven by a generally active promoter expressed throughout the retina, and *Nrl-dsRed* restricted to the outer INL and ONL where rods reside (**Fig. 5o**). Furthermore, we found that cell death was distributed in the INL and GCL (**Fig. 5n, o**, blue) as shown for normal retina at this stage (**Fig. 1f-g**). When we compared the number of dying cells in the rod lineage population (dsRed+) that received the control *sgCtl-Cas9-eGFP* plasmid with those that did not, there was no difference (**Fig. 5p**, left). However, when we quantified cell death in those rod lineage cells (dsRed+) that received the experimental *sgGrik3-Cas9-eGFP* plasmid and controls, there was a significant difference with *Grik3* targeting resulting in low levels of rod lineage cell death (**Fig. 5p**, right). This indicates that GRIK3 functions to promote the death of developing rod photoreceptors.

## Human retinal development shows hallmarks of an OPN4-ipRGC-rod pathway

Collectively, data from analysis in the mouse indicate that rod photoreceptor precursors are pruned from the retina in response to light activation of OPN4 in ipRGCs. To determine whether this pathway was conserved during human retinal development, we performed several analyses.

First, we took an *in silico* approach to estimate the human gestational age that was equivalent to the period in the mouse (P0-P7) when the OPN4-ipRGC-rod precursor pathway is functioning. Previous studies employed open-ended dynamic timewarping (OE-DTW) on bulk retinal transcriptomes from human and mouse development to align the developmental time-courses^35^. Employing a similar analysis, we find that the period of OPN4-ipRGC-rod precursor pathway function corresponds to the mid-2^nd^ trimester human retinal age (Day (D) 100 - Day 136+; **Fig. 6a**).

**Figure 6:**
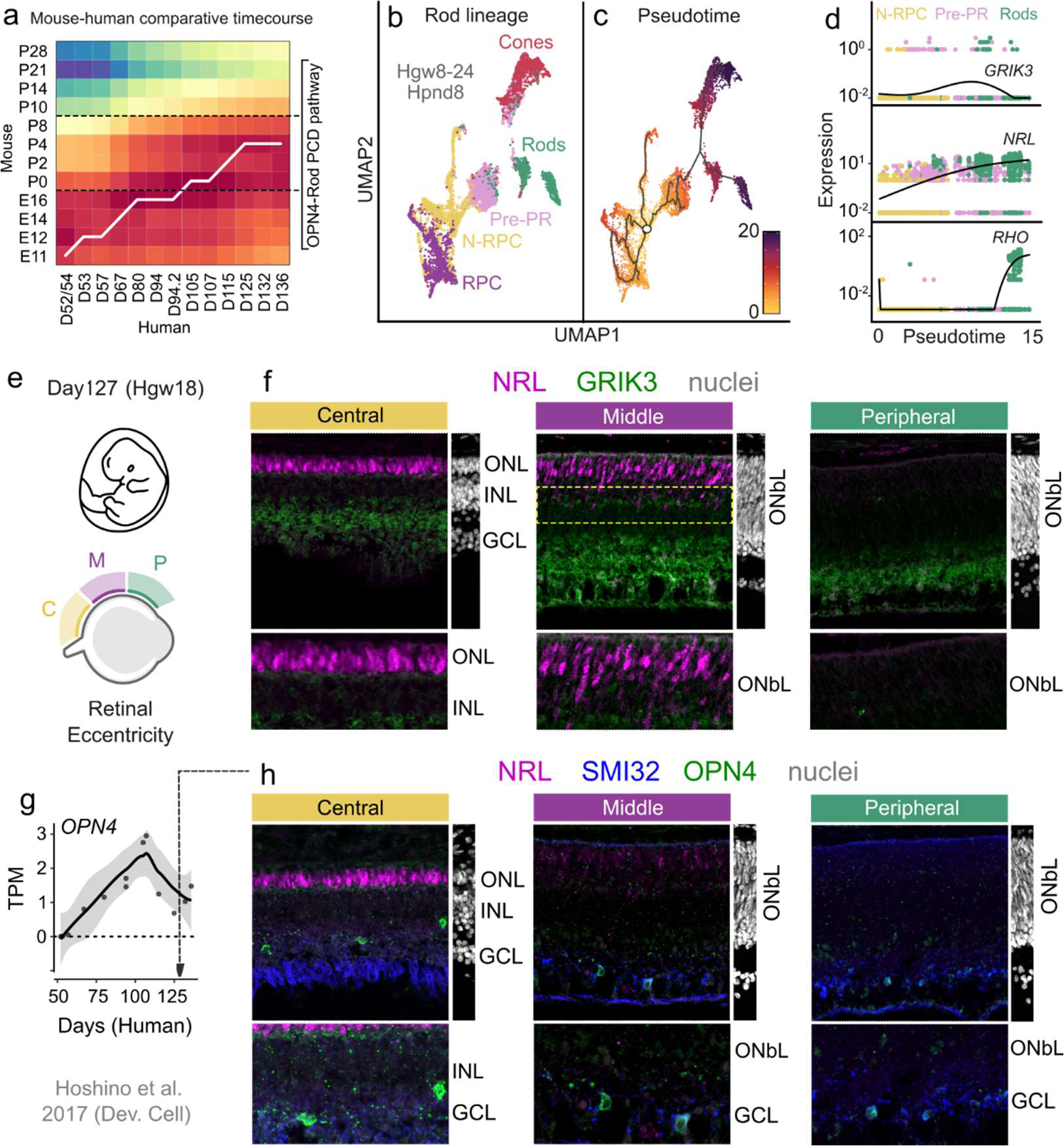
Hallmarks of an ipRGC-to-immature rod axis during human fetal development. **a**, Bulk transcriptome alignment of developing mouse and fetal human retina (Hoshino et al., 2017) using open-ended dynamic timewarping (OE-DTW), displayed as a heatmap with line of best-fit between samples, D = gestational day, E = embryonic day, P = postnatal day. **b**, scRNA-seq profiling and UMAP embedding of human fetal retinal development from Lu et al., 2020. **c**, pseudotime analysis from neurogenic retinal progenitors (N-RPCs) to developing rods; Hgw = human gestational week, Hpnd = human postnatal day. **d**, Analysis of developmental expression of *GRIK3* (top), *NRL* (middle), and *RHO* (bottom) in pseudotime in **b**. **e**, Schematic representation of spatio-temporal development of the human retina, capturing a temporal axis along eccentricity: P = peripheral, M = middle, C = central retina. **f**, Representative images of NRL+ and GRIK3+ cells from Hgw18 human retina across regions. **g**, Bulk transcriptomic profiling of human retinal development from Hoshino et al., 2017 (as in **a**), higlighting temporal expression of *OPN4*. **H,** *In vivo* histological assessment of fetal human ipRGCs using a primate-specific OPN4 antibody and immature rods via NRL, characterized over eccentricity. GCL = ganglion cell layer, IPL = inner plexiform layer, INL = inner nuclear layer, NBL = neuroblastic layer, OPL = outer plexiform layer, ONL = outer nuclear layer, ONbL = outer neuroblastic layer, TPM = tags per million. Regression in **G** is represented as mean ± s.e.m as the shaded area.

We next surveyed fetal human development for transcriptional indicators of the OPN4-ipRGC-rod precursor pathway. Mining a scRNA-Seq atlas of human retinal development^36^ (**Fig. 6b**), we performed pseudotime analysis of the developing rod lineage, akin to earlier mouse analyses. We then projected the expression levels of three human homologues, *RHO*, *NRL* and *GRIK3* that are known to be markers of the mouse OPN4-ipRGC-rod precursor pathway onto the human pseudotime axis (**Fig. 6d**). This revealed a dynamic expression pattern very similar to that observed in the mouse (**Fig. 5**) with transient *GRIK3* peaking as *NRL* expression is rising prior to *RHO* expression (Figure 6d). Furthermore, analysis of human fetal retina (gestational week 18) by immunofluorescence shows that NRL and GRIK3 are distributed in a manner similar to the mouse (**Fig. 6e, f**). Human retinal development occurs with a central to peripheral progression gradient such that peripheral is the least developed and central retina the most mature^35^. This means that it is possible to examine multiple developmental stages within a specimen. In the D127 (Hgw18) example shown, limited NRL labeling is observed in the least developed peripheral retina (**Fig. 6f**, right). However, in central D127 retina, NRL is observed in rods at the outer edge of the neuroblastic layer (ONbL) as well as within the displaced, immature rods that are precursors (**Fig. 6f**, center, yellow dashed box). At this stage, GRIK3 labeling is also apparent in the band corresponding to the rod precursors (**Fig. 6f**, central, yellow dashed box). Finally, we assessed expression of OPN4. In bulk transcriptome analysis of human retina^35^, *OPN4* is expressed at detectable levels with peak expression at about D107 (**Fig. 6g**). This developmental stage corresponds precisely to the stage of mouse development (P0) when the OPN4-ipRGC-rod begins to function.

Assessment of retinal sections at ages following the peak transcript expression revealed cells in the GCL and INL that were immunoreactive for human hOPN4. Despite suboptimal preservation of human tissue samples, we also observed lines of hOPN4 labeled puncta suggesting the existence of neurites laminating in the inner and outer IPL (**Fig. 6h**, see **Extended Data Fig S5a** for mouse). Like NRL expression, hOPN4 was highly expressed in cells within the central retina with lower expression towards the periphery (**Fig. 6h**). Importantly, we find several examples of OPN4+ punctate structures leaving the IPL and projecting towards the outer retina. This suggests conservation of outer retinal processes in human fetal ipRGCs. The combination of these data suggest that hallmarks of an ipRGC-to-immature rod axis are conserved during human retinal development.

## Discussion

Here we reveal that early light stimulation of OPN4-expressing ipRGCs leads to pruning of immature rods through a transient, retrograde pathway within the retina. This pathway involves a subset of photosensitive ipRGCs that extend neurites into the outer retina where they interdigitate with rod photoreceptor precursors. The pruning of rod photoreceptors is light-dependent and according to deletion of *Slc17a6/Vglut2* in ipRGCs and of the glutamate receptor gene *Grik3* in rod precursors, is mediated by the neurotransmitter glutamate. In mice, this pathway is active in the early postnatal period prior to eye-opening, an indication that this pathway is developmental and is in preparation for, rather than a component of, visual experience.

During early retinal development, many neuron classes (RGCs, amacrine cells, bipolar cells) are overproduced and pruned away through intrinsic and extrinsic mechanisms^37^. While prior work has focused on how inner retinal neuron apoptosis occurs^38^ and its role in visual behavior^39^, outer retinal photoreceptor pruning has remained relatively unexplored. Early studies surveying naturally occurring developmental apoptosis have noted immature rod apoptosis within the INL that peaks towards the end of the first postnatal week (∼P7/P8)^17,40,41^. However, unlike inner retinal neurons that primarily utilize Bax-dependent mechanisms for programmed cell death, immature rods require both Bax and Bak^42^. Consequently, combined Bax and Bak loss-of-function leads to decreases in apoptosis of rod photoreceptors within the developing and mature INL^42^. In the current study, we establish that developmental cell death of the immature rod population depends on the lighting environment and occurs non-autonomously. Importantly, this appears to be selective information channel that routes sensory information to developing rod photoreceptors in the INL and not to other potential cell classes (**Fig. 2e**). This selective channel requires ipRGC-input through melanopsin-based photodetection (OPN4) and ipRGC glutamate packaging/release (VGLUT2), being detected autonomously by developing rods that transiently express *Grik3*.

It was previously shown that ipRGCs extend neurites into the outer retina^8,18,19^. These structures were originally termed outer retinal dendrites (ORDs) but based on the analysis performed here they appear to have the anatomic characteristics of dendrites, but molecular features of axons. More specifically, in addition to expression of the OPN4 photopigment suggesting local photodetection, they contain the glutamate packaging protein VGLUT2 and the presynaptic marker synaptophysin. They may thus be best thought of as a hybrid neurite. The current analysis shows that in the mouse, outer stratifying ipRGC processes are abundant at the day-of-birth but diminish in number over the first postnatal week, consistent with the period in which immature rod cell death is regulated.

Our analysis also suggests that an OPN4-ipRGC-immature rod pathway might be conserved and active during human fetal development. By assessing pseudo-temporal gene expression^36^ of *GRIK3* in the rod photoreceptor lineage and histologically analyzing immature rod (NRL+ cells) lamination in Hgw18 retina, we find that these cells transiently make their way into the developing INL, as observed in the mouse. Furthermore, this occurs during a period in human development where OPN4 is expressed in retinal ganglion cells (*OPN4* transcript and OPN4 protein), some of which are noted to have OPN4+ puncta projecting towards the developing outer retina. Thus, even with ∼90 million years separating humans and mice, it appears that hallmarks of this developmental pathway remain conserved. As such, an OPN4-ipRGC-immature rod pathway in human development has several implications.

First, the developmental staging would suggest that photoreception by ipRGCs in utero may regulate rod photoreceptor numbers. Prior investigation of mouse retinal development and OPN4-signaling has shown that ipRGCs are responsive to light during late gestation (E16)^43^, a stage that corresponds to ∼Day 80-94 in the human (**Fig. 6a**). Consistent with this timing, a human epidemiological study found an association between day length during early gestation and the risk of severe retinopathy of prematurity (ROP)^44^ and is thought to be dependent on OPN4 in the mouse^6^. Finally, in infants born very preterm, it has been noted that outer retinal photoreceptor responses are less sensitive and robust than term matched controls^45,46^. This suggests that at stages of viable premature birth (∼Day 150 and beyond), the aberrant lighting environment experienced by these newborns may result in inadvertent immature rod elimination via an ipRGC-to-rod developmental pathway.

## Materials and Methods

### Sample size estimation and investigator blinding

For all analysis and experiments presented, sample sizes were not predetermined. However, for all experiments comparing metrics between genotypes, lighting conditions, etc. investigators were blinded during the experiment. When this was not possible, unbiased methods such as machine-learning assisted cell counting (cellpose 1.0^47^) were applied so that investigator input could not bias the analysis.

### Mice

Mice of both sexes were housed in a pathogen-free vivarium, maintained at 22 °C. All viral and surgical procedures were conducted in accordance with protocols approved by the Institutional Animal Care and Use Committee at Cincinnati Children’s Hospital Medical Center. Genetically modified mice used in this study include: *Opn4*^KO^ (^48^), *Opn4*^cre^ (^49^), *Opn5*^cre^ (^7^); *Ai14* (^50,51^), *INTACT* (^16^), *Morf3* (^21^), *Rx^cre^* (^52^), *Th*^flox^ (^53^), *Nrl^cre^* (^15^), *Vglut2^flox^*(^29^), *Drd4^fl^* (this study).

### Human fetal sample acquisition

Human eyes we recovered from donors with no identifiers between post-conception D84 (Hgw12) and D127 (Hgw18) under an approved protocol through the University of Washington (UW5R24HD000836). The age of human specimens were estimated using a combination of clinical metrics: crown-rump length, gestational ultrasound, fetal foot length measurements when possible. In total eyes from the following ages were used for this study: D84 (1 donor), D85 (1 donor), D89 (1 donor), D96 (1 donor), D98 (2 donors), D127 (1 donor).

### Environmental lighting and housing

Mice were housed in standard vivarium fluorescent lighting (photon flux 1.6 × 10^15^ photons cm^−2^ sec^−1^) supplemented with violet light emitting diodes (4.9 × 10^14^ photons cm^−2^ sec^−1^ in the 375-435 nm range). Mice were housed in a 12h:12h light:dark cycle unless otherwise noted. Dark-rearing experiments were conducted by moving the pregnant dam to a light-tight chamber the night before giving birth. For the duration of the experiment, only dim red LEDs were used for cage changes or handling.

### Intravitreal injections for ipRGC trafficking

P0-1 pups were anesthetized on ice for 4 minutes before being placed on a pre-chilled ice pack under a stereo-microscope. A lateral incision was made at the presumptive eyelid suture using a gauge 22 needle. A pilot incision was made into the ora serrata using the tip of a gauge 29½ micro-syringe, followed by viral delivery into the vitreous using a pulled glass micropipette attached to a Drummond II Nanoinjector. 300nL of AAV/DJ-hSyn1-DIO-tdTomato-T2A-Syp::eGFP-WPRE (2.5 × 10^13^ GC/mL from Addgene #51509) was injected unilaterally at a rate of 60nL/sec. The tip of the micropipette was left in the vitreous for 15 seconds after the final pump to prevent backflow. Following surgeries, pups were placed under a heat lamp until they recovered and reintroduced to the dam. 6-7 days following intravitreal injections, mice were sacrificed and retina were harvested for imaging. Retina with damage from needle perforation were excluded from the analysis.

For post-eye opening injections, mice were anesthetized via inhaled isoflurane (4% induction, 2% maintenance). A gauge 27 needle was used to make a pilot incision into the ora serrata, followed by insertion of a 10uL Hamilton syringe loaded with 1.5uL of AAV/DJ-hSyn1-DIO-tdTomato-T2A-Syp::eGFP-WPRE (2.5 × 10^13^ GC/mL from Addgene #51509). Following intravitreal delivery, the syringe remained in the eye for 15 seconds to limit backflow. Mice were allowed to recover from anesthetic on a heating pad, returned to their home cage when ambulatory, and assessed twice a day until they were sacrificed 7 days later.

### Histology and Tissue Processing

*Tissue harvest and processing.* Retinal whole mounts and brain sections were processed as previously described. Briefly, mice were anesthetized via lethal isoflurane inhalation followed by decapitation (P0-P12) or cervical dislocation (P12+). Eyes were enucleated and placed into 4% paraformaldehyde (PFA) for 45 minutes at room temperature. Brains were processed similarly but allowed to incubate in fixative overnight at 4C on an orbital shaker. Retinae were isolated from the fixed eye and stored in 1x phosphate buffered saline (PBS) at 4C until further processing. For cryosectioning and staining, retina were dehydrated in 15% sucrose for 15 minutes, followed by 30% sucrose treatment overnight. The tissue was then embedded in OCT medium, frozen on dry ice, and sectioned at 12-16um. For analyses comparing conditions and genotypes, to limit staining variability, multiple retina from each condition was embedded in the same OCT block, sectioned, and stained together.

*Immunostaining and Histology*. Tissue sectioning and staining were performed as previously described. Retina and brain sections were permeabilized in 0.5% Triton-X in PBS (PBST), followed by blocking using 10% normal donkey serum in 0.5% PBST. Tissues were stained with primary antibody overnight at room temperature, washed five times with 1x PBS and incubated with secondary antibodies for 1h at room temperature. Subsequently, sections were washed five more times with 1x PBS, coated in mounting media (Vectashield) and coverslipped for imaging. Serial sections of retina were acquired for staining and histology, all regions of the retina were sampled for this analysis. For wholemount immunostaining, retinas were permeabilized in 0.5% PBST for 2h at room temperature, followed by 2h of blocking. Incubation with primary antibodies occurred for 2-3 days at 4C on an orbital shaker. The retinas were washed for 30 minutes with 1x PBS, this was repeated 6-8 times, followed by secondary antibody incubation at room temperature for 1 hour on an orbital shaker. Then, retinas were washed with 1x PBS 6-8 times, after which they were flattened and coverslipped. Primary antibodies used in this analysis include:

### Single ipRGC reconstruction via MORF3-labeling

High resolution z-stack images of ipRGCs from *Opn4^cre/+^; Morf3^+/-^; Ai14^+/-^* were obtained using a 40x objective on a Nikon AXR Confocal, with a z-step of 0.3µm. Z-stacks comprised the entire depth of the retina for pre-eye opened ages and the inner retina for post-eye opened ages. For each time point we captured P1: 27 fields (45 cells), P5: 18 fields (30 cells), and P18: 23 fields (30 cells). These images were imported into FIJI (v1.3.5) and reconstructed semi-automatically using the SimpleNeuriteTracer plugin (SNT) in 3D. SWC files were saved and imported into R and reconstructions in XY, XZ, and YZ dimensions were generated using the natverse library.

### Fos activation assessment of ipRGCs and targets

Mice (pregnant dams for P0 experiments, P4 pups for P5 experiments) were dark adapted at least 6 hours prior to the experiment in light-tight chambers. Pups were placed on a heating pad set to 37C and illuminated with 1×10^14^ photons cm^−2^ sec^−1^ light using a narrowband LED centered at 470 nm. Control mice were placed on an identical heating pad in complete darkness. After 90 minutes of illumination, pups were sacrificed and tissue was immediately harvested, fixed in 4% PFA as described, and processed for analysis. Retina were serially sectioned and random slides were chosen for Fos topological analyses, with ∼5-8 sections analyzed per animal. Brains were serially sectioned and slices corresponding to the IGL (P0) or SCN (P5) were assessed for Fos induction.

### Bulk RNA-sequencing of developing retinal tissue

*Tissue harvesting and library* generation. *Opn4^+/-^* × *Opn4^+/-^* harem crosses were used to produce wildtype and knockout littermate pups raised in similar lighting environments (∼250 lux). On the day of birth (P0), tail and fingerclippings were harvested and mice were genotyped prior to determine appropriate genotypes. At P3, mice were sacrificed between ZT6-7 and retina were dissected in ice-cold oxygenated Ames’ media (95% air, 5% oxygen). Multiple retinas were pooled together from animals of the same genotype (4 retina per sample, 2 animals), and tissue was snap frozen in Trizol (100uL). Post-hoc genotyping was then performed on each sample to confirm genotypes. RNA extraction, library preparation, and sequencing (Illumina HiSeq 2 × 150bp) were all performed at GeneWiz. Samples with >500ng of RNA and a RIN > 8.5 were selected for further processing. The average read depth per sample was 42.7 ± 3.9 million reads. Subsequently, adapters and low-quality reads were trimmed/filtered using Timmomatic v0.39. Reads were pseudoaligned to the mouse reference genome (GRCm38) and counts were quantified using Kallisto v0.46.1. Differential gene expression analysis was performed using Sleuth v0.30.8.

*CoGAPS and Transfer Learning* via projectR. Tags per million (TPM) output from Kallisto of 254 differentially expressed genes from each sample were used for CoGAPS and transfer learning. Briefly, this subset expression matrix (gene × sample) was converted into two weighted CoGAPS patterns (WT or KO enriched) using the setParam(“nPatterns”, 2) and CoGAPS() functions in the CoGAPS package. To project the featureLoadings of the weighted patterns into single cell latent space, we first chose to subset the Clark et al. 2019 atlas to 10,000 cells with the top 3000 variable genes, and then projected featureLoadings from the weighted patterns onto the scRNA-Seq dataset using the projectR() function with default parameters.

*Bulk Deconvolution of bulk RNA-Seq.* The gene count matrix from Kallisto was used for single-cell reference mapping and bulk deconvolution. First the bulk count matrix was converted into an ExpressionSet object in R using the Biobase::ExpressionSet() function from the Biobase library. The Clark et al. 2019 atlas was processed as follows: genes expressed in <3 cells were excluded from analysis and the dataset was scaled, normalized, and mitochondrial gene expression regressed using the SCTransfrom() function in the Seurat v4 package. Using the publication-provided metadata, we assigned each cell to a class, and determined the markers of each class using the FindAllMarkers() function with only.pos = T, min.pct = 0.25, and logfc.threshold = 0.25. Marker genes and cell proportions from the scRNA-seq atlas were used to generate a pseudobulk dataset using the BisqueRNA::ReferenceBasedDecomposition() function with non-overlapping markers. Then, for each bulk RNA-seq sample, we de-convolved and inferred the cell class proportion using the BisqueRNA::MarkerBasedDecomposition() function with weighting = True. Importantly, we used only cells captured between P0 and P8 from the scRNA-seq dataset as this best represents the time points of our bulk analysis.

### EdU pulse chase and proliferation estimation

EdU (5-ethylnyl-2’-deoxyuridine) was prepared according to the manufacturer’s recommendations (10mM stock solution in DMSO, ThermoFisher #C10337). This solution was then diluted in sterile PBS to achieve a final concentration of 5mg/kg body weight of the pup when injected. EdU was administered subcutaneously and allowed to incorporate into proliferating cells for 1 hour as described in **S Fig 1A**. Mice were sacrificed, retinae were harvested, and EdU+ cells were analyzed per animal (*n =* 4 / genotype / age). For innate proliferation in *Opn4* WT and *Opn4* KO littermates, retinae were harvested at P5, sectioned and stained against markers of proliferation (pH3, Ki67) and progenitor cells (Vsx2 low). Then, images corresponding to half the section (hemi-section) were acquired using a 10x objective. To linearize the retina, a spline was fit to the OLM and the image was transformed using the spline as an anchor. Following image processing, cells corresponding to triple positive were assessed and quantified by a blinded investigator.

### Spatial analysis of neuronal processes, cell location, and apoptosis

To assess and standardize the spatial location of ipRGC sprouts, soma location, and dying cells, we devised a method to spatially normalize data to the outer limiting membrane (OLM) or inner plexiform layer (IPL). Hemi- or whole retinal section were imaged using either a 10x or 20x objective and imported into FIJI for analysis. To generate unified spatial metrics to assess sprouts, cell death, and spatial locations, we first generated maximum intensity projections of each image and used channels with highest background fluorescence to determine OLM/IPL positions. Using the ‘segmented line’ tool, a line was manually fit to cognate structure and a ‘Straighten..’ function was applied to linearize the curved tissue. Then, using the multipoint tool, points were added along the newly transformed image’s OLM/OPL to extract [X,Y] coordinates for normalization and distance metrics.

IpRGC sprout tips were located manually by a trained investigator and using the multipoint tool, their locations were noted, exported, and normalized to IPL-Y coordinates. To unbiasedly detect CC3+ soma over hemi- and whole retinal sections, we used a generalist convolutional neural network (CNN) – cellpose 1.0^47^ to automatically detect cells. Masks for each cell were then imported in FIJI^47^. These masks were then overlaid on channels containing signals from cell type markers for further characterization. Positional information from these masks were exported and normalized to OLM-Y coordinates for spatial analysis.

### Rod spatial analysis and 2D nearest neighbor estimation

One year old *Opn4*^WT^ and *Opn4*^KO^ littermates were reared in light-controlled chambers and used for rod number estimation and spatial analysis. Retina were harvested and serially sectioned at 10um as described, tissue was stained against Nrl (to mark rod nuclei), cone arrestin 3 (Arr3; labeling all cones) and DNA to label all cells in the outer retina. 16 fields of outer retina were acquired per animal (*n* = 4 per genotype) and >21.5k cells and their positions were acquired and normalized to the OPL. For analysis, per field estimates of rod and cone number were used from each animal. For nearest neighbor calculations, within each field, rod nuclei centroid coordinates were computed and then the nndist() function was applied using the spatstat library.

### Pan RGC-development scRNA-seq datamining

Single cell transcriptomes and associated metadata from E14-P5 sorted RGCs were acquired from the Gene Expression Omnibus (GEO) under GSE185671 and https://github.com/shekharlab/mouseRGCdev. To subset developing M1 ipRGCs (c33 and c40 from Tran et al., 2019), we used provided Waddington optimal transport (WOT) metrics precomputed where cells from the developing scRNA-seq dataset were considered M1s if their combined M1-WOT score was ≥ 95^th^ percentile across all ages. This yielded a total of 4,280 putative M1s over development (E14: 949, E16: 826, P0: 1226, P5: 1279 cells). Gene_x_cell matrices corresponding to these cells were processed using the Seurat v4 pipeline stated previously, with the exception of integration to correct batch effects using 2000 variable features. The integrated object was scaled, transformed, and the first 30 principle components were used for clustering using default parameters. For each cluster, we used the FindAllMarkers() function to determine if any clusters comprised cells that were non-M1s. These clusters were subsequently excluded from analysis.

### In vivo co-electroporation and somatic CRISPR mutagenesis

To somatically knockout *Grik3* from developing rods, we generated small guide RNAs to PAM sites in proximity to exons 4 and 7 of the murine *Grik3* locus and cloned these into the pSpCas9(BB)-2A-GFP (PX458) backbone (Addgene #48138), where expression of the sgRNAs is controlled under the U6 promoter. Co-electroporation and labeling of rods were accomplished by expressing of dsRed under the Nrl promoter (*p*Nrl), and was achieved using the *p*Nrl-dsRed plasmid (Addgene #12764). Both constructs were purified at a concentration of ∼4ug/uL and were prepared as a 1:1 mixture for electroporation. 300uL of solution was injected using a pulled glass microneedle into the subretinal space between P0-2. Then, using a 10mm tweezerode (Harvard Apparatus, Inc #45-0119) 5 square-wave pulses were applied at 80V for 50ms with an inter-stimulus interval of 950ms. Mice were then toe- and tail-clipped, allowed to recover under a heating lamp and returned to the dam after all surgeries were complete. After 5-8 days, electroporated retina were harvested as described and analyzed for EGFP and dsRed expression histologically. Exclusion criteria included direct damage to the electroporation site, low EGFP and/or dsRed expression. This led to only a small fraction of mice (9/112, 0.8%) having strong signal-to-noise for each reporter and little to no damage to the electroporated area for analyses.

### Human fetal development – Transcriptomic and histological analysis

*Open-Ended Dynamic Time Warping*. To align mouse and human retinal development transcriptomes to determine cross-species age estimates, we employed open-ended dynamic time warping (OE-DTW) as previously described for human/mouse alignment in the retina. First, multiple mouse samples representing the same time point were averaged and used for OE-DTW. Previous work established 3,072 dynamically expressed genes during human retinal development (), which were then one-to-one orthologue matched to mouse using the HOM_MouseHumanSequence.rpt file obtained from Mouse Genome Informatics (). This yielded 2,667 conserved genes between human and mice whose expression was then used for OE-DTW analysis. Dynamic time warping was performed using the dtw package (v) in R using the dtw() function with arguments for open.end = TRUE and dist.method = “Manhattan”.

*Tissue Histology.* Following extraction of the eye, the anterior chamber was punctured and the entire globe was fixed in 4% PFA for 45 minutes. Following this, the lens was removed, and the eye cup was allowed to further fix in 1% PFA + 3% sucrose solution overnight. The cup was washed 3 times with 5% sucrose in PBS and then incubated overnight in the following sucrose solutions in PBS. 10% sucrose, 20% sucrose, 30% sucrose, 50% sucrose, and finally a 30% sucrose in PBS:OCT mixture. Then eyecups were transferred to 100% OCT and snap frozen in ethanol with dry ice. Tissue sections were then cut on a cryostat at 20um thickness and stained using identical protocols to that used for mouse histology and immunofluorescence.

*Spatial analysis of human retinal features*. Similar to mouse analysis, retinal tissue was processed for imaging and image analysis. Whole section images of the DAPI channel plus any other channels were first acquired using a 4x objective and tiling. Then, the section was divided into three equally spaced regions from the optic disc, giving rise to peripheral, middle, and central regions in our analyses. Then, higher magnification images were acquired using a 20x objective for any spatial or marker-based analysis. For spatial normalization of datapoints, images were spatially transformed to linearize the retinal section (using a spline function in ImageJ), then the OLM-centric coordinates were used for all analyses to normalize to a retinal landmark.

### Statistical Analysis

All statistical analyses were performed in RStudio and R v4.0.5 using base stats or the rstatix package. Statistical tests and corresponding sample sizes (*n*) are noted in the figure legends. For all analyses utilizing multiple comparisons, or multiple independent tests of a hypothesis, *P*-values were corrected via false discovery rate correction (fdr correction in text). Sample sizes were not predetermined.

## Data Availability

Bulk transcriptomic data generated in this study will be uploaded to GEO under Accession # (XXXX) and be made available upon paper publication.

## Acknowledgements

We thank Paul Speeg (Lang lab) for excellent mouse colony management. We would also like to thank Gowri Nayak, PhD and Yueh-Chiang Hu, PhD of the CCHMC Transgenic Animal and Genome Editing Core Facility for generating genetically modified mouse lines, Grik3-targeting plasmids, etc. We thank Shiming Chen, PhD (Washington University in St. Louis) for the Nrl-cre transgenic mouse line and Anand Swaroop, PhD (NIH National Eye Institute) for the BP-cre mouse line. This work was supported by NIH grants R01 EY027077, R01 EY027711, R01 EY032029, funds from the Goldman Chair of the Abrahamson Pediatric Eye Institute at CCHMC to RAL, T32NS007453 training grant to KG, R01 EY025555 and P30EY003039 to PG, Damon Runyon Cancer Research Foundation grant DRG-# 32-20 and Hanna H. Gray Fellows Program Award (HHMI) GT15994 to KCE, and the Albert J. Ryan Predoctoral Fellowship to SPD.

## Author Contributions

SPD and RAL conceived the project. SPD designed and executed experiments. SPD and BAU performed transcriptomic and expression-related computational analysis. KCE and IG acquired human fetal tissue for histological analysis. MTN performed initial cell death assessments. KG and SPD performed viral tracing experiments. AA and SPD performed neurite spatial analyses.PG generated key primate-specific OPN4 antibodies. SPD analyzed the data. SPD and RAL wrote the manuscript.

