## Extended Data Fig. for "Developmental adaptation of rod photoreceptor number via photoreception in melanopsin (OPN4) retinal ganglion cells"

### Extended Data Figures & Legends:

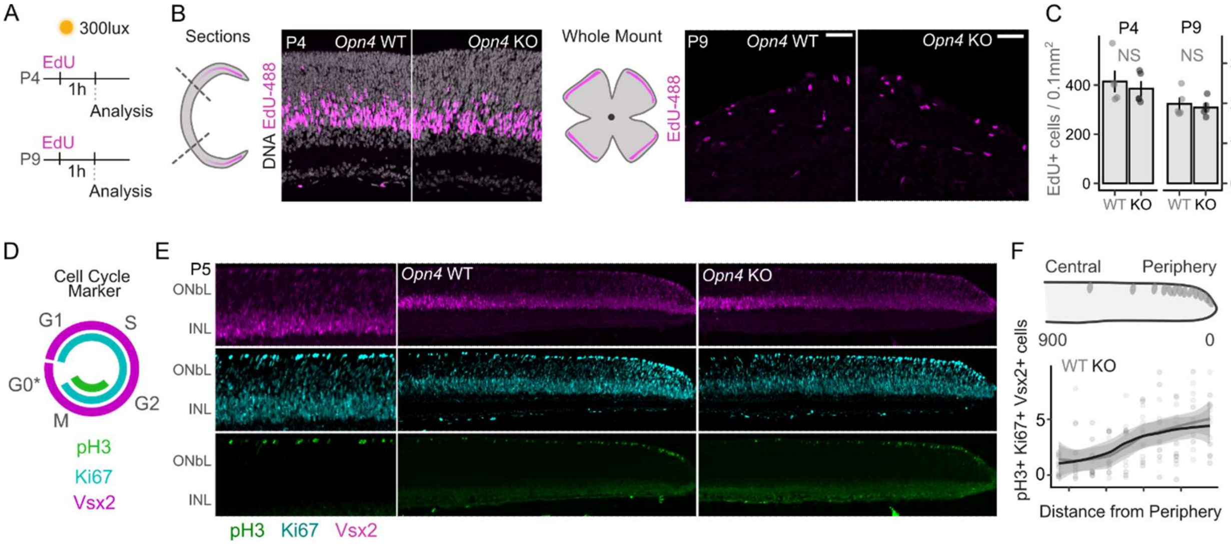


**Figure S1 | ipRGC phototransduction does not influence proliferation in the retina**

**A**, Schematic of EdU (5-ethynyl-2'-deoxyuridine) pulse chase experiment performed on P4 and P9 *Opn4*$^{WT}$ and *Opn4*$^{KO}$ littermates. **B**, Representative images from sectioned (left) or whole-mount (right) retina following pulse chase from each genotype. **C**, Analysis of proliferating cell numbers from images in **B** (NS = not significant, 4 mice per genotype per age). **D**, Schematic representation of cell cycle marker expression in the retina. **E**, Laminar localization of cells expressing pH3, Ki67, and Vsx2 in *Opn4*$^{WT}$ and *Opn4*$^{KO}$ mice (*n* = 6 per genotype). **F**, Spatial quantification of triple positive cells (pH3+ Ki67+ Vsx2+) across eccentricity reveals no influence of ipRGC phototransduction on proliferation at P5. GCL = ganglion cell layer, IPL = inner plexiform layer, INL = inner nuclear layer, NBL = neuroblastic layer, OPL = outer plexiform layer, ONL = outer nuclear layer.

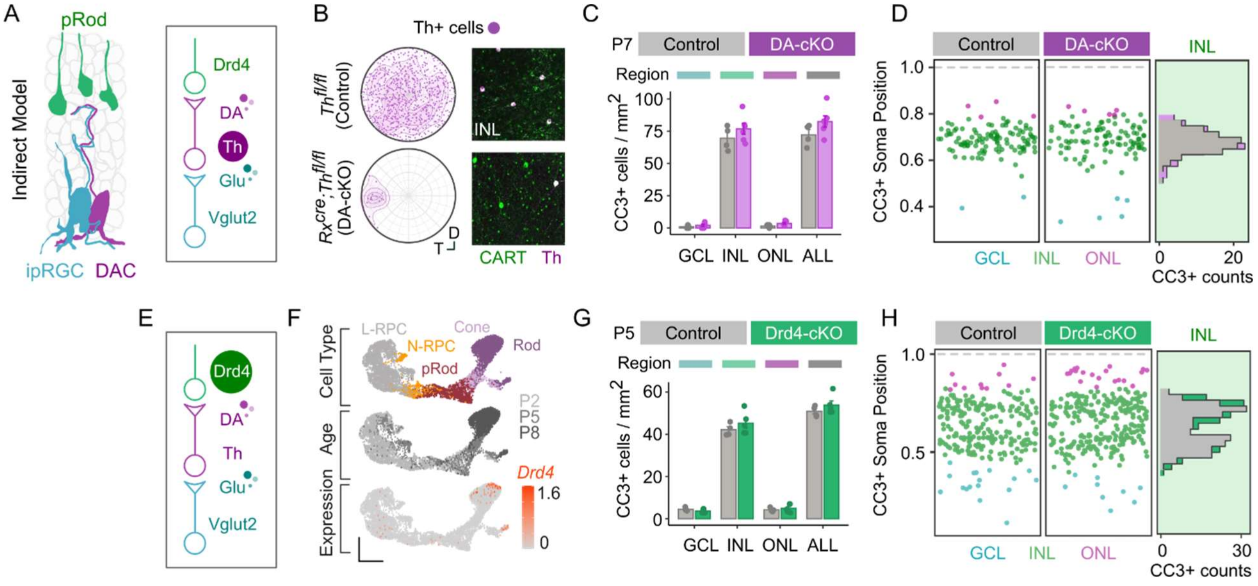


**Figure S2 | Dopamine signaling does not contribute to retinal apoptosis during development**

**A**, Schematic representing intra-retinal circuit linking glutamate release to rods via a dopaminergic amacrine cell axis. **B**, Assessment of Th loss in control (*Th*$^{fl/fl}$) and DA-cKO (*Rx*$^{cre}$*; Th*$^{fl/fl}$) mice stained against Th and Cartpt (marker of dopaminergic amacrine cells). **C**, Spatial analysis of retinal cell death during development from control and DA-cKO mice at P7 showing limited changes across quantity and space. **E**, Circuit diagram testing Drd4-mediated dopamine sensing as a putative mechanism of sensory-evoked rod apoptosis. **F**, tSNE embedding of rod lineage from Clark et al., 2019 highlighting expression of Drd4 late in rod development (detectable only at P8 in scRNA-seq). **G**, Spatial analysis of retinal cell death at P5 from control (*Drd4*$^{fl/fl}$ ) and Drd4-cKO (*Rx*$^{cre}$*; Drd4*$^{fl/fl}$) retina showing a limited contribution of Drd4 signaling in rod apoptosis during development. GCL = ganglion cell layer, IPL = inner plexiform layer, INL = inner nuclear layer, NBL = neuroblastic layer, OPL = outer plexiform layer, ONL = outer nuclear layer. Data is represented as mean ± s.e.m.

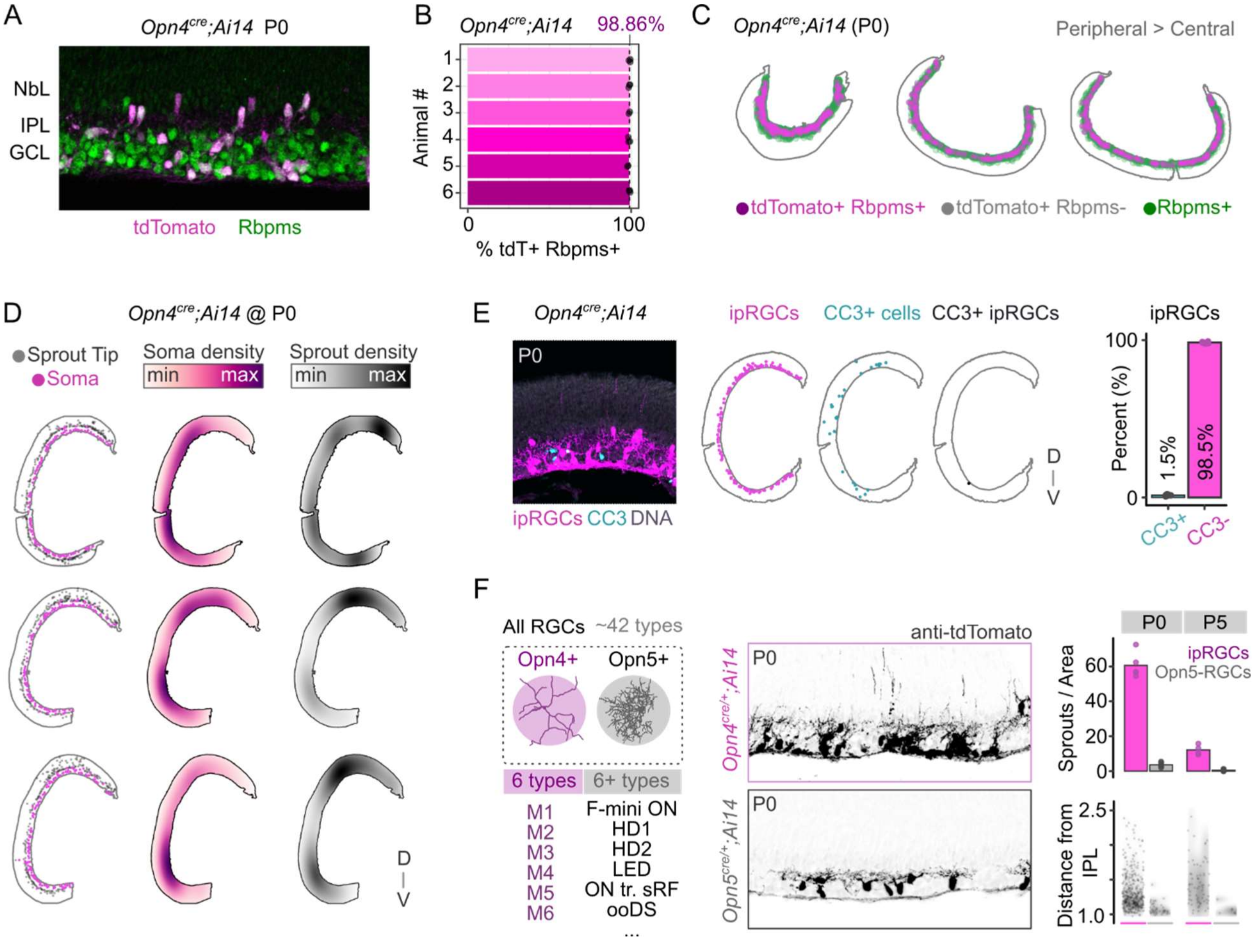


**Figure S3 | Characteristics of lineage-marked and sprout-bearing cells in the *Opn4$^{cre/+}$; Ai14* mouse line across retinal development**

**A-C,** Analysis of tdTomato expression specificity to retinal ganglion cells (RGCs, Rbpms+) at P0, quantifying overlap (**B**) and spatial distribution (**C**) across *n* = 6 mice. **D**, Spatial analysis of ipRGC soma position and sprout location across retinal sections, highlighting point distributions (left), soma densities (middle), and sprout densities (right). **E**, Histological analysis of ipRGC apoptosis (left), quantification of spatial locations of ipRGCs, CC3+ cells, and CC3+ ipRGCs (middle), and analysis of percentage of the ipRGC population actively dying (CC3+) at P0 (right). **F**, Left, Distinction of ipRGC types from Opn5-RGC types. Middle, Representative images from P0 *Opn4$^{cre/+}$; Ai14* and *Opn5$^{cre/+}$; Ai14* retina showing tdTomato+ RGCs in the ganglion cell layer. Right, quantification of sprout number and sprout tip distance from IPL between ipRGC and Opn5-RGCs. Data for ipRGCs is repeated from **Figure 3**.

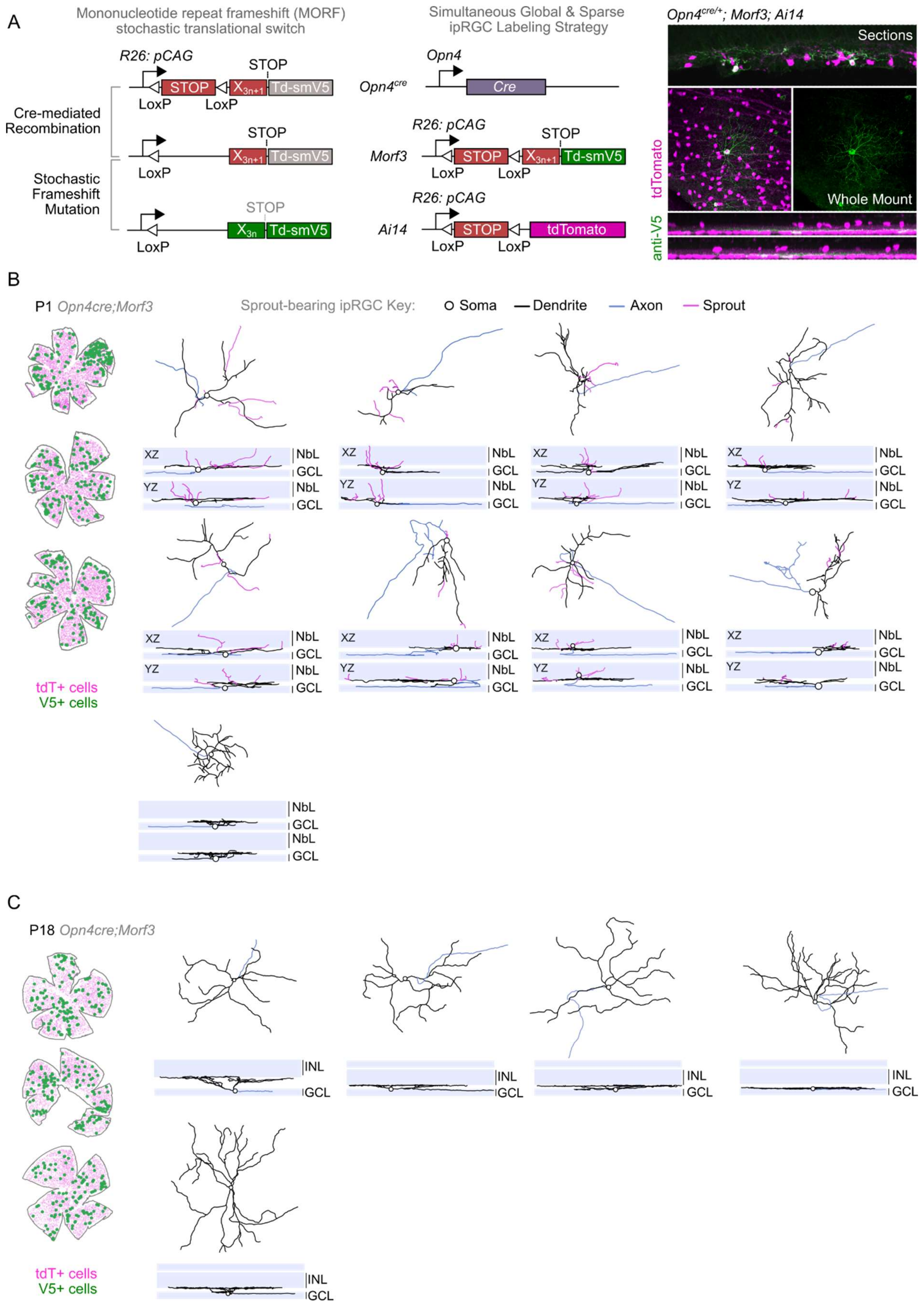

A
Mononucleotide repeat frameshift (MORF) stochastic translational switch
R26: pCAG
STOP
LoxP
LoxP
Td-smV5
Cre-mediated Recombination
Stochastic Frameshift Mutation
Simultaneous Global & Sparse ipRGC Labeling Strategy
Opn4
Opn4cre
Cre
Morf3
Ai14
tdTomato
Opn4cre/+; Morf3; Ai14
Sections
Whole Mount
anti-V5
B
P1 Opn4cre;Morf3
Sprout-bearing ipRGC Key:
Soma
Dendrite
Axon
Sprout
XZ
YZ
NbL
GCL
tdT+ cells
V5+ cells
C
P18 Opn4cre;Morf3
INL
GCL
tdT+ cells
V5+ cells

**Figure S4 | Single ipRGC reconstructions via MORF3 reporter**

**A,** Left, Schematic highlighting the Morf3 reporter expression system, which requires the expression of cre-recombinase and a random mono-nucleotide frameshift in the Morf3 allele to allow for appropriate translation of a membrane-bound V5 epitope tag in cells. Middle, Lineage tracing framework used to label developing ipRGCs sparsely and globally by generating *Opn4*$^{cre/+}$*; Morf3*$^{+/-}$*; Ai14* mice with representative images located towards the right. **B**, Example whole retinal soma maps of all ipRGCs (magenta; tdT+) and sparsely labeled ipRGCs (green, V5+). Right, several examples of sprout-bearing ipRGCs (#1-8) and non-sprout bearing cells (#9) projected en-face, or orthogonally (XZ, YZ). Additionally, cells #6 and #8 bear both axon collaterals and sprouts. **C**, Similar to B but with ipRGCs reconstructed post eye-opening (P18). GCL = ganglion cell layer, IPL = inner plexiform layer, INL = inner nuclear layer, NBL = neuroblastic layer, OPL = outer plexiform layer, ONL = outer nuclear layer.

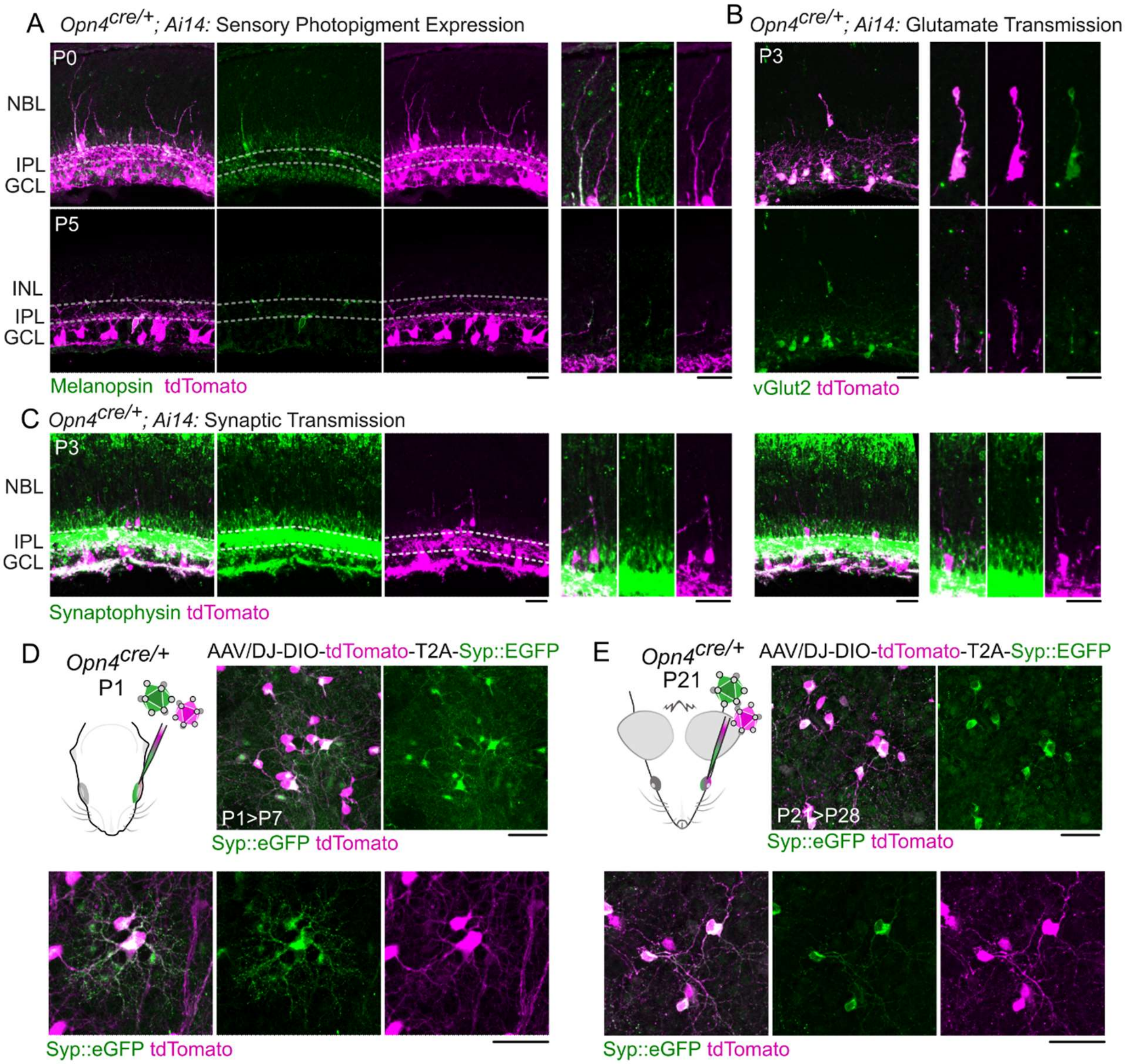


**Figure S5 | Molecular fingerprinting of ipRGC dendritic sprouts during development**

**A,** Expression of melanopsin within sprouts in *Opn4cre; Ai14* mice at P0 and P5. **B**, Expression of Vglut2 in ipRGC sprouts at P3. **C**, Expression of native synaptophysin (Syp) in the retina of *Opn4$^{cre}$; Ai14* mice at P3, highlighting expression in ipRGC sprouts but higher expression in other cells (rods, cones, amacrine cells). **D**, Intravitreal injection of AAV-DIO-tdTomato-T2A-Syp::EGFP prior to eye opening highlighting strong dendritic expression in ipRGCs. **E**, Similar to E but injected post eye-opening (P21) to label older ipRGCs at P28. GCL = ganglion cell layer, IPL = inner plexiform layer, INL = inner nuclear layer, NBL = neuroblastic layer, OPL = outer plexiform layer, ONL = outer nuclear layer.

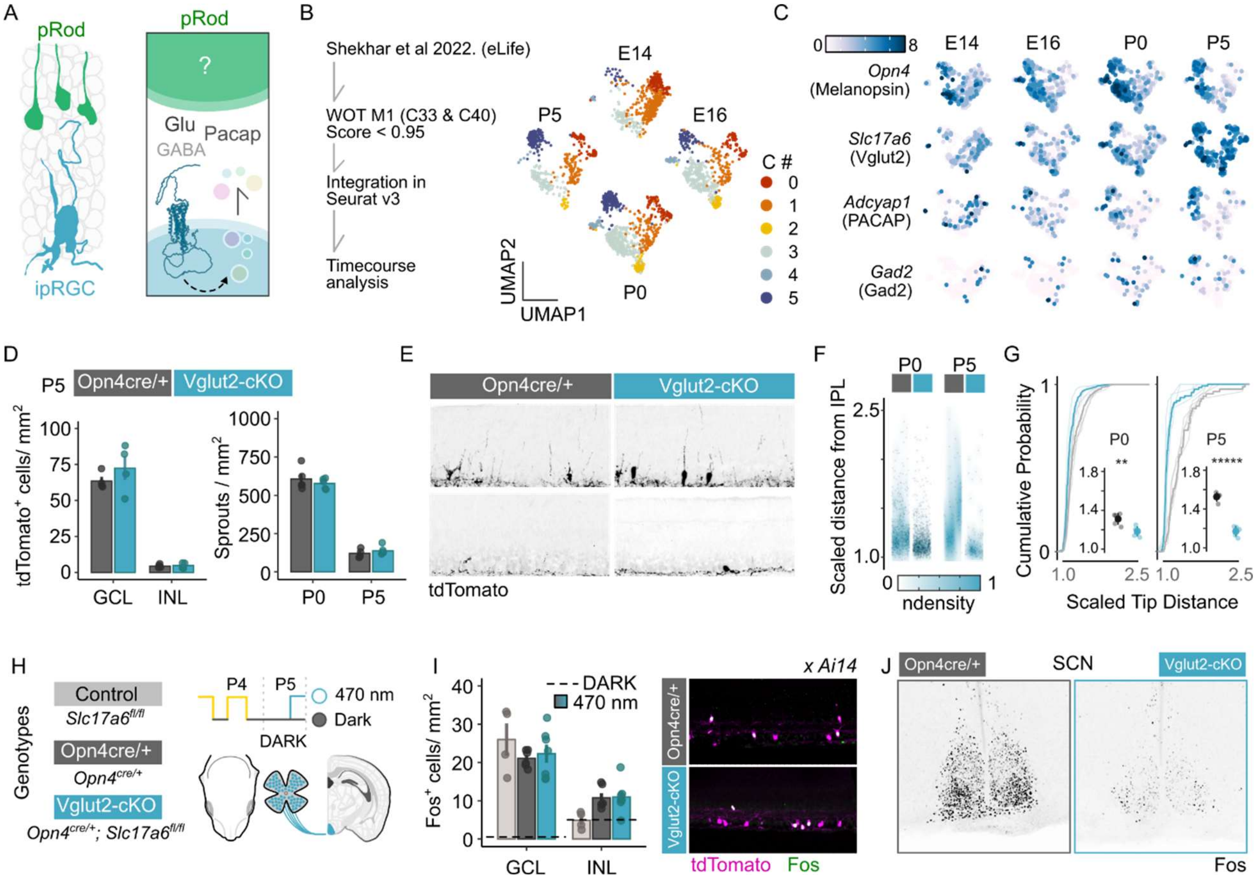


**Figure S6 | Conditional loss of Vglut2 from ipRGCs leads to minimal autonomous alterations.**

**A**, Schematic representing various modes of ipRGC neurotransmission. **B**, Left, Analysis pipeline utilizing scRNA-seq profiling of murine RGCs across fetal and postnatal development (Shekhar et al, 2022, see methods for details). Right, UMAP embedding of clusters representing developing M1 ipRGCs (C33, C40) from E14 – P5. **C**, Normalized expression of melanopsin (*Opn4*), Vglut2 (*Slc17a6*), PACAP (*Adcyap1*) and Gad2 (*Gad2*) during development of M1 ipRGCs. **D**, Comparison of cell numbers and sprout numbers between *Opn4$^{cre/+}$; Ai14* controls and Vglut2-cKO (*Opn4cre/+; Slc17a6$^{fl/fl}$; Ai14*) highlighting no cell loss or sprout loss. **E**, Representative images from **D**. **F-G**, Analysis of scaled sprout laminar distance from the IPL (IPL = 1) between control and Vglut2-cKO mice across development with average scaled tip distance quantified ($n$ = 5 mice per genotype, $^{**}P < 0.01$ and $^{*****}P < 0.000001$, Students $t$ test with fdr correction), P0 differences in absolute length: Opn4creAi14 vs Opn4cre;Vglut2fl/fl;Ai14 = 12.07um (95% CI: 2.64 21.51), P5 differences in absolute length: Opn4creAi14 vs Opn4cre;Vglut2fl/fl;Ai14 = 20.29um (95% CI: 13.33 27.25. **H**, Fos induction experiment, similar to Fig 1J, comparing activation of ipRGCs and post-synaptic targets in the brain. **I**, Quantification and representative images of Fos+ cells from all three genotypes reveals no difference in light sensitivity to 1 x $10^{14}$ photons $cm^{-2}$ $sec^{-1}$ 470 nm stimulation (NS = not significant, Tukey HSD with fdr correction comparing genotypes within regions). **J** Representative images of the SCN from light-stimulated control and Vglut2-cKO mice. GCL = ganglion cell layer, IPL = inner plexiform layer, INL = inner nuclear layer, NBL = neuroblastic layer, OPL = outer plexiform layer, ONL = outer nuclear layer. Data is represented as mean ± s.e.m.
